# Intron retention of an adhesion GPCR generates single transmembrane-helix isoforms to enable 7TM-adhesion GPCR function

**DOI:** 10.1101/2023.01.11.521585

**Authors:** Anne Bormann, Marek B. Körner, Anne-Kristin Dahse, Marie S. Gläser, Johanna Irmer, Vera Lede, Judith Alenfelder, Joris Lehmann, Daniella C.N. Hall, Michael Thane, Michael Schleyer, Evi Kostenis, Torsten Schöneberg, Marina Bigl, Tobias Langenhan, Dmitrij Ljaschenko, Nicole Scholz

## Abstract

Adhesion G protein-coupled receptors (aGPCR) function as metabotropic mechanosensors in the nervous system and other organs. aGPCR are heavily spliced forecasting an extraordinary molecular structural diversity. Many predicted isoforms lack the transmembrane (7TM) signaling subunit, but to what extent these non-GPCR isoforms are produced and what physiological purpose they serve is unknown. Alternative splicing through intron retention of *ADGRL/Latrophilin/Cirl* mRNA in *Drosophila* generates transcripts encoding unconventional proteins with an extracellular domain anchored by a single transmembrane helix (Cirl^1TM^). Here, we show that *Cirl^1TM^* transcripts are translated *in vivo* and that Cirl^1TM^ binds Cirl^7TM^ N-terminal fragment-dependently. This interaction enables mechanosensory neurons to distinguish input intensities through Gα_o_-dependent signaling. Similarly, a direct interaction was found for mammalian GPR126/ADGRG6 isoforms. Together, our findings define intron retention and isoform-specific heteromerization as extraordinary molecular strategies to adjust *Cirl*-dependent mechanosensation and demonstrate physiological relevance of versatile aGPCR isoform repertoire to tune cellular responsiveness.

## INTRODUCTION

Mechanosensing is vital for nervous system development, morphology and function^1–4^. Disturbances of the delicate mechanobiological system of nervous tissue are pathologically relevant and linked to traumatic brain injury, multiple sclerosis and brain cancer^2,5–8^. Unfortunately, thus far, detailed knowledge about the molecular mechanisms at play remains scarce.

Cells of the nervous system can express several dozen G protein-coupled receptors (GPCR)^9^. Their signature layout with an extracellular N terminus, intracellular C terminus and a seven-transmembrane helix-spanning domain (ECD, ICD, 7TM domain respectively) is well-established and considered basic to transduce extracellular into intracellular signals. This way, GPCR assume roles in fundamental neurobiological processes in neurons and glia^10,11^. On the pathological flip side, GPCR are considered one of *the* pharmaceutical targets to treat neurological disorders^12^. In recent years, adhesion-type GPCR (aGPCR), a class within the GPCR superfamily, have emerged as metabotropic mechanosensors, relevant in several organ systems including the nervous system^13–19^. For example, in zebrafish, rapid electrical signal propagation depends on ADGRG6/GPR126 as it times the onset of peripheral neuron myelination likely by probing the matrix rigidity at the neuron-Schwann cell interface^13,20^. In *Drosophila*, sensitivity towards sound, touch and proprioceptive cues hinges on ADGRL/Latrophilin/Cirl function^21,22^. In mammals, vision and hearing are associated with ADGRV1/VLGR1 mediating mechanical stretch response through its association with focal adhesion contacts^23–26^.

aGPCR are encoded by large genomic loci and are known to be heavily alternatively spliced^27,28^. However, there are only few examples where isoform-specific functions of aGPCR have been uncovered. For example, the S4 variant of ADGRG1/GPR56 with its shortened N-terminal fragment (NTF) was shown to be a vital regulator in synaptic pruning^29,30^; an N-terminally truncated ADGRL3/LPHN3 isoform shapes the transmission of cone photoreceptor synapses ^31^. Outside of the nervous system, different ADGRL3/LPHN3 isoforms (with a varying loop region) decrease insulin secretion from isolated pancreatic β cells^32^. Recent bioinformatic efforts helped to systematically catalog the aGPCR transcriptome^27^. Fascinatingly, ~ 22% of all aGPCR genes produce transcripts that lack the sequence encoding a complete 7TM domain^27^, rendering G protein coupling to 7TM-less receptor isoforms highly unlikely. This begs questions as to whether such transcript variants are translated into protein, especially *in vivo*, how their expression varies across time and space, if they contribute to physiologically relevant cellular functions, and if these extend beyond an adhesive role. To address these questions, we studied the ADGRL/Latrophilin homolog *Cirl* (Ca^2+^-independent receptor of latrotoxin) from *Drosophila*. *Cirl* is predicted to undergo alternative splicing to generate transcripts encoding peculiar non-GPCR proteins with only NTF, the first transmembrane helix and an alternative C terminus (Cirl^1TM^). Cirl was shown to be relevant in mechanosensing. It suppresses cAMP levels and likely modulates gating properties of ionotropic mechanosensors in proprioceptive sensory neurons of the lateral pentascolopidial chordotonal (lch5) organ, a well-defined set of cells accessible to genetic, optical and functional approaches^21,22^.

Here, we report that *Cirl^1TM^* transcripts are produced *in vivo*, through intron retention and that they are co-expressed with *Cirl^7TM^* transcripts throughout development. Contradictory to previous accounts suggesting degradation and/or physiological dispensability of intron retention-based molecules^33^, we demonstrate the expression of the non-GPCR protein Cirl^1TM^ in a living animal. Moreover, we show that Cirl^1TM^ forms a heteromeric complex with Cirl^7TM^ through its NTF. Strikingly, our functional analyses provide evidence that these molecular Cirl^1TM^-Cirl^7TM^ assemblies define the discriminative bandwidth of proprioceptive sensory neurons towards mechanical stimulation, a function conferred through Gα_o_ protein-dependent signaling. Importantly, we documented a similar complex formation for mammalian ADGRG6^7TM^ and unconventional ADGRG6^1TM^ proteins.

In sum, our findings uncover that alternative splicing of aGPCR enables cooperation between GPCR-like and non-GPCR isoforms to adjust cellular mechanophysiology providing a promising entry point for experimental and therapeutic approaches to interfere with aGPCR signals.

## RESULTS

### Intron retention produces *Cirl* transcripts encoding molecules that contain a single TM domain and alternative intracellular region

In *Drosophila*, Cirl/Latrophilin/ADGRL is encoded by a single gene^22^. Alternative splice events of *Cirl* pre-mRNA produce eight different transcripts (flybase.org) putatively encoding six unique proteins (Figures 1A, S1A,B). The majority of transcripts encode classical GPCR that contain a 7TM domain allowing G protein engagement, β-arrestin binding and canonical intracellular signaling (Cirl^7TM^ = *Cirl-G, -I, -B, -E, -H*; Figures 1A, S1A,E). However, a substantial fraction of transcripts encode proteins containing only the N terminus integrated in the plasma membrane through a single TM helix (Cirl^1TM^ = *Cirl-J, -F, -K*; Figures 1A, S1B,E). *Cirl^1TM^* transcripts likely arise through intron retention, where a specific splice donor is ignored and intron 6 is retained. The resulting transcripts are either degraded or translated. In the latter case, translation would continue until an intronic in-frame stop codon is reached producing 207 alternative amino acid residues of the putative Cirl^1TM^ isoforms (Figure 1A).

**Figure 1.**
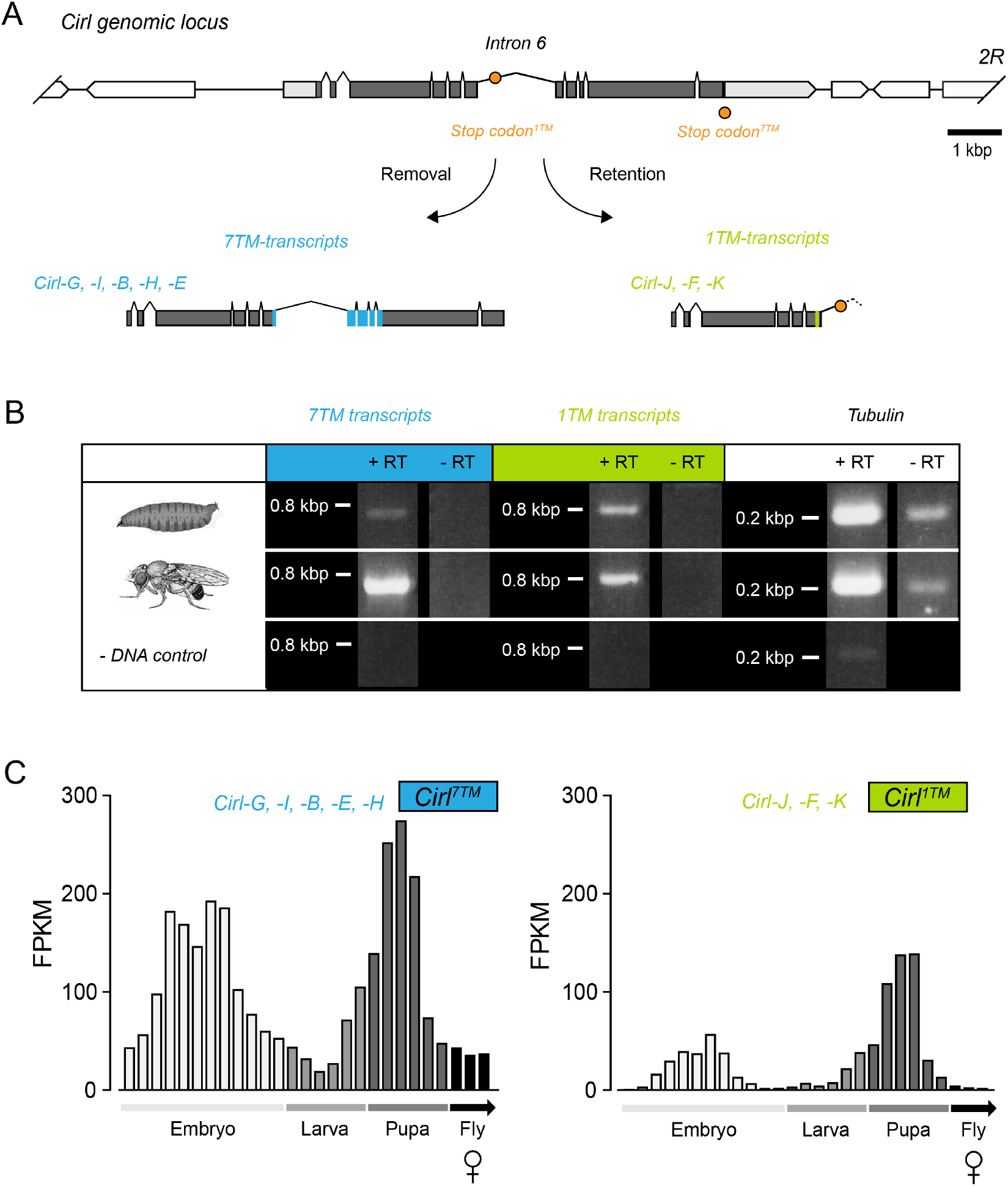
Intron retention produces *Cirl* transcripts encoding molecules with a single TM and alternative intracellular region. **(A)** Schematic of the *Cirl* gene locus. Removal of intron 6 gives rise to five *Cirl^7TM^* transcripts: *Cirl-G, -I, -B, -E*, and *-H* (in blue). Retention of intron 6 generates three *Cirl^1TM^* transcripts: *Cirl-J, -F, -K* (in green). Depicted are only protein coding portions of the transcripts. Resulting proteins are shown in Suppl. Figure S1. **(B)** Validation of *Cirl^1TM^* transcript in *Drosophila*. RT-PCR products amplified from larval and adult cDNAs for *Cirl^7TM^* (blue column; 0.7 kbp) and *Cirl^1TM^* transcripts (green column; 0.8 kbp). Tubulin served as loading and quality control of cDNA and RT-PCR; no DNA: specificity control. **(C)** Cumulative fragments per kilobase million (FPKM) values of *Cirl^7TM^* (left panel) and *Cirl^1TM^* (right panel) at several developmental time points. While *Cirl^7TM^* and *Cirl^1TM^* have a similar ontogenetic expression profile, *Cirl^7TM^* transcript quantity was higher than that of *Cirl^1TM^* throughout development. Embryo (white): 12 time points equally distributed between 0 and 24 h; larval stages (light gray) L1, L2 and L3 (split across 4 time points) in days; pupa (dark gray): 6 time points across 4 days; female flies (black): 1, 5, 30 days old. The individual transcript quantity is given in Suppl. Figure S1.

To test for the presence of *Cirl^1TM^* transcripts, we used *Cirl^1TM^* and *Cirl^7TM^*-specific primer sets to conduct RT-PCR analyses on cDNA libraries generated from adult and larval *w^1118^ Drosophila*. We amplified *Cirl^1TM^* and *Cirl^7TM^*-specific DNA fragments (Figure 1A,B), implying co-expression of both in the developing and adult fly. To exclude the possibility that small amounts of yet to be degraded *Cirl^1TM^* transcripts served as PCR template and to investigate whether *Cirl^1TM^* and *Cirl^7TM^* are consistently co-expressed, we quantified the amount of each *Cirl* transcript generated at consecutive time points during embryonic, larval, pupal, and adult development employing an RNA-sequencing data set (Figure S1C,D;^34^). *Cirl^7TM^* and *Cirl^1TM^* transcripts show similar ontogenetic expression profiles with peak abundances during embryonic and pupal stages (Figures 1C, S1C,D). *Cirl^1TM^* and *Cirl^7TM^* account for ~ 22.5% and ~ 77.5% of total *Cirl* transcript produced, respectively (Figure S1E). These data validate the expression of *Cirl^1TM^* transcripts and suggest that *Cirl^1TM^* and *Cirl^7TM^* transcript expression is co-regulated. This implies the need of both transcript species during specific developmental periods.

### Cirl^1TM^ protein is expressed *in vivo*

To ascertain if Cirl^1TM^ protein is expressed *in vivo*, we used the previously generated *Cirl^KO^-attP* allele to knock-in *Cirl* variants that encode Cirl^1TM^ isoforms containing V5-tags at the C terminus and Cirl^7TM^ isoforms containing Flag-tags in the third intracellular loop, thus enabling their specific detection through matching antibodies. In addition, a monomeric RFP (mRFP) in the NTF of Cirl (Figure 2A,B)^21^ was introduced to detect the entire isoform repertoire. This allele is referred to as *Cirl^3x-tagged^*. Since RNA-sequencing data suggested elevated expression of *Cirl^1TM^* during pupal stages (Figures 1C, S1C,D), pupal protein extracts were analyzed in Western blots using a monoclonal α-V5 antibody to detect a ~33 kDa fragment corresponding to the C-terminal fragment (CTF) of Cirl^1TM^ (Figure 2C).

**Figure 2.**
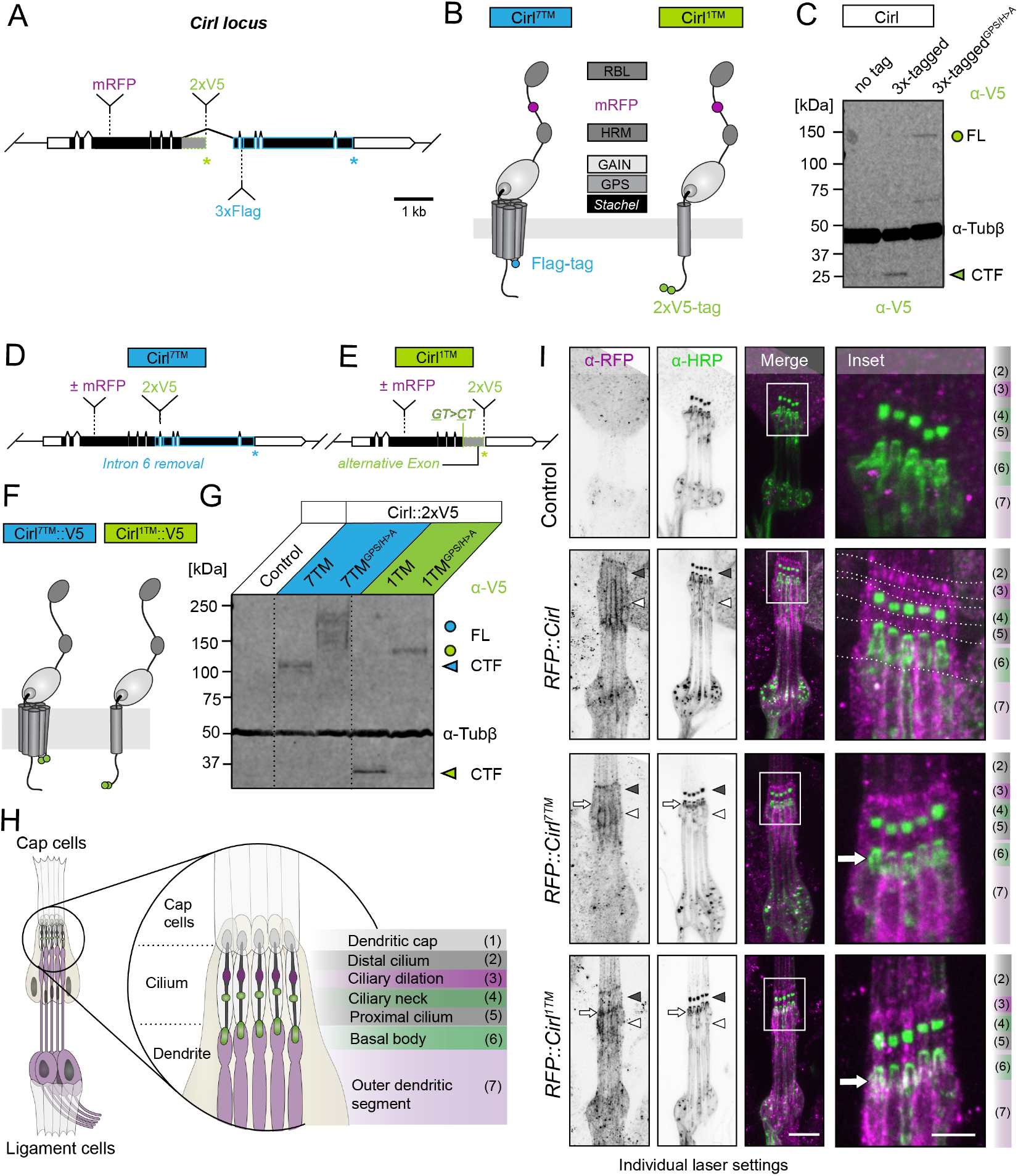
Cirl^1TM^ protein is expressed in mechanosensory neurons. **(A)** Schematic of *Cirl* locus. Insertion sites of an mRFP (indicated in magenta), a tandem V5 tag (2xV5, in green) and a triple Flag tag (3xFlag, in blue) are indicated. **(B)** Tagging strategy of Cirl^7TM^ and Cirl^1TM^ proteins. mRFP was inserted in the linker region between RBL and HRM domain^21^. Tandem V5 tag is located at the C terminus of Cirl^1TM^, triple Flag tag is included in the third intracellular loop of Cirl^7TM^. **(C)** Western blot of lysates from pupae, which express V5-tagged Cirl^1TM^ protein. Bands at ~ 33 kDa (green arrowhead) and ~ 136 kDa (green circle) correspond to the expected molecular weight of CTF and FL Cirl^1TM^, respectively. Additional band at ~ 70 kDa of unknown identity. Tubulin served as a loading control (α-Tub*β*). Related to Figure S2. **(D)** Schematic layout of *Cirl* allele exclusively expressing Cirl^7TM^. Intron 6 encoding Cirl^1TM^-C-terminus was removed. **(E)** Schematic layout of *Cirl* allele exclusively expressing Cirl^1TM^. Splice donor at the exon 6-intron 6 boundary was mutated (<u>G</u>T > <u>C</u>T) and 7TM-specific exons and introns were deleted. **(F)** Schematic of Cirl^7TM^ and Cirl^1TM^ proteins fused to 2xV5 in the third intracellular loop (left) and the very C-terminal end (right), respectively. **(G)** Western blot analysis of head lysates from Cirl^7TM^∷V5 or Cirl^1TM^∷V5 showed CTFs with molecular mass of ~ 110 kDa and 33 kDa, respectively. Cleavage-deficient versions with predicted molecular masses of ~ 190 kDa (FL Cirl^7TM-GPS/H>A^∷V5) and ~136 kDa (FL Cirl^1TM-GPS/H>A^∷V5). *Cirl^Rescue^* = untagged control. Tubulin served as a loading control (α-Tub*β*). **(H)** Schematic illustration of lch5 organ with 5 bipolar sensory neurons and typical anatomical features, numbered (1) – (7). **(I)** Subcellular localization of Cirl isoform species in the lch5 organ (intensity inverted α-RFP and α-HRP channels are shown in black and white). Maximum z-projections of confocal images show RFP∷Cirl^7TM^ at the level of the cilium (gray arrowheads), in the ciliary dilation (3), outer dendritic segments [(7), white arrowheads]. RFP∷Cirl^1TM^ location overlaps with RFP∷Cirl^7TM^ at similar subcellular region. In addition RFP∷Cirl^1TM^ is located at the basal bodies [(6), indicated by arrow]. *Cirl^Rescue^* = Control (upper row). α-HRP recognizes neuronal membranes and was used to outline the lch5. Scale bar 10 μm, inset: 5 μm. Related to Figure S3.

GPCR autoproteolysis-inducing (GAIN) domain-dependent autoproteolytic processing presents one of the hallmark features of aGPCR and underlies their bipartite NTF/CTF structure. Autoproteolysis of Cirl can be averted by mutating the GPCR proteolysis site (GPS)^21^, at which nascent receptors are self-cleaved. To further confirm the expression of Cirl^1TM^ proteins, we generated a cleavage-deficient version of *Cirl^3x-tagged^* via a −2 GPS H>A mutation (*Cirl^3x-tagged/H>A^*) and recovered full-length (FL) Cirl^1TM^ protein (~137 kDa; Figure 2C). Consistent with this finding, Western blot analysis of head homogenates from *Cirl^3x-tagged^* and *Cirl^3x-taggedH>A^* animals showed Cirl-NTF (~110 kDa), FL Cirl^1TM^ (~135 kDa) and FL Cirl^7TM^ (~215 kDa; Figure S2) expression also in the central nervous system of adult flies. Thus, *Cirl^1TM^* transcripts are translated to produce Cirl^1TM^ proteins *in vivo*.

### Cirl^1TM^ protein localizes to mechanosensory neurons

Previously, we reported on Cirl’s role in mechanosensitive lch5 neurons^21,22^ but the *Cirl^KO^* allele used in those studies removes both Cirl^7TM^ and Cirl^1TM^ isoforms precluding analyses of their individual contributions. Consequently, we set out to decipher isoform-specific functions of Cirl, especially Cirl^1TM^ proteins.

To gain genetic control over Cirl isoform expression, we engineered *Cirl* alleles for exclusive expression of either Cirl^1TM^ or Cirl^7TM^ gene products under endogenous transcriptional control (Figure 2D,E). Exclusive Cirl^7TM^ expression was assured by removing the intron (intron 6 of isoform *-I* and intron 5 of *-E, -B/H, -G*) which otherwise could encode the Cirl^1TM^-specific alternative intracellular tail (Figure 2D). Exclusive Cirl^1TM^ expression was achieved by a silent mutation of the splice donor (*<u>G</u>T* → *<u>C</u>T*) at the exon 6/intron 6 boundary and removal of the *Cirl^7TM^*-specific genetic region (Figure 2E). To validate this genetic strategy, we conducted Western blot analyses from head homogenates of flies in which we inserted biochemical V5 tags at the C terminus of Cirl^1TM^ (wild-type: Cirl^1TM^∷V5; cleavage-deficient: Cirl^1TM/H>A^∷V5; Figure 2F,G) and the third intracellular loop Cirl^7TM^ proteins (wild-type: Cirl^7TM^∷V5; cleavage-deficient: Cirl^7TM/H>A^∷V5; Figure 2F,G). We detected the expected FL and CTF bands of both Cirl^1TM^∷V5 and Cirl^7TM^∷V5 fusion proteins (Figure 2G).

α-V5 immunofluorescent detection via confocal imaging of Cirl^1TM^∷V5 in lch5 organs failed due to high signal background (data not shown). To locate Cirl^1TM^ in lch5 organs, we generated an independent set of Cirl^1TM^ or Cirl^7TM^ alleles carrying an additional mRFP within the NTF (referred to as RFP∷Cirl^1TM^∷V5 and RFP∷Cirl^7TM^∷V5)^21^ (Figure S3). α-RFP immunolabeling and microscopy of larval lch5 organs revealed specific RFP∷Cirl^1TM^∷V5 and RFP∷Cirl^7TM^∷V5 signals at the level of the ciliary and outer dendritic segments as well as at the ciliary dilation (Figure 2H,I), demonstrating that both Cirl isoform species co-localize in these subcellular compartments. Moreover, in the outer dendritic segment, particularly around the basal body, we detected mostly RFP∷Cirl^1TM^∷V5 (Figure 2I) showing exclusive localization of this isoform. Thus, mechanosensory lch5 neurons express both Cirl^7TM^ and Cirl^1TM^ proteins.

### Cirl^1TM^ and Cirl^7TM^ proteins are co-required for neuronal discrimination of mechanical inputs

*Drosophila* larvae rely on *Cirl-*dependent proprioceptive feedback from lch5 function for locomotion (Figure 3A,B)^22,35^. This begs the question whether Cirl^1TM^, Cirl^7TM^ or both isoform species are involved in this behavior. Quantification of the average midpoint crawling speed of L3 larvae showed that sole expression of Cirl^7TM^ suffices to revert values back to those of the *Cirl^Rescue^*. Interestingly, exclusive presence of Cirl^1TM^ even enhances movement velocity above *Cirl^Rescue^* values (Figure 3A,B, Table S3). While this may implicate that Cirl^7TM^ suppresses Cirl^1TM^ function relevant for this particular behavior, coordinated locomotion results from numerous neurons and neuronal processes, some of which may require Cirl. To pinpoint the functional relevance of different Cirl isoform species, we performed electrophysiological recordings from mechano-stimulated lch5 neurons. Previously, measurements from lch5 neurons showed that *Cirl^KO^* larvae discriminate less robustly between different vibrational stimulation frequencies than *Cirl^Rescue^*^21,22^. The used direct vibration stimulation lacked control of stimulus amplitude^21,22^. To increase stimulation precision in lch5 recordings and to emulate proprioceptive stretch-induced lch5 activity more adequately, we attached a miniature hook to a piezo element and pulled at the organ using defined solitary mechanical square pulses (Figure 3C,D). The hook was rapidly moved by defined distances between 0.03 and 3 μm perpendicularly to the lch5 organ. This way the organ is stretched triggering action current (AC) frequencies with remarkably low variability.

**Figure 3.**
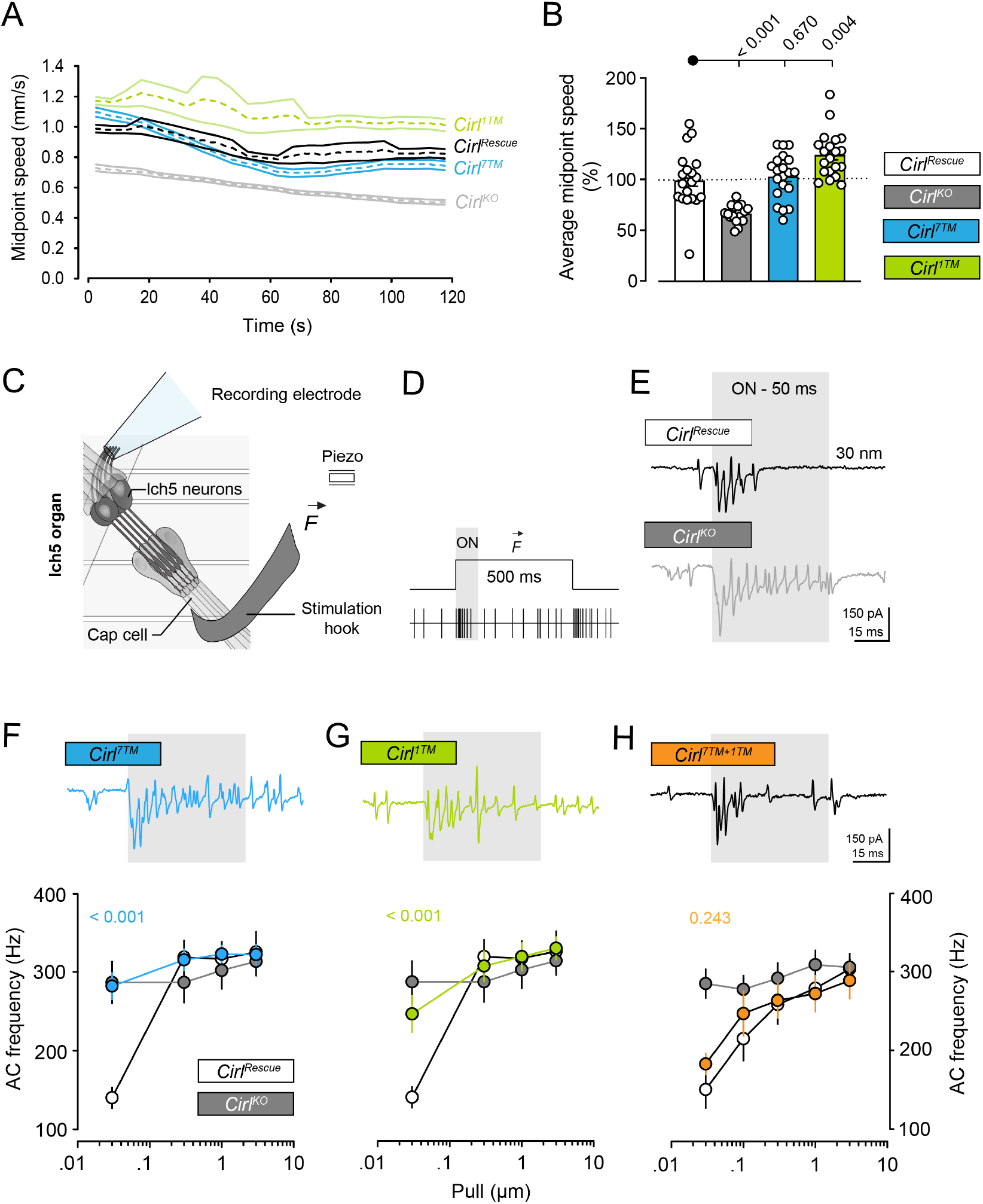
Cirl^1TM^ and Cirl^7TM^ proteins are co-required for neuronal discrimination of mechanical inputs. **(A)** Groups of *Cirl^KO^*, *Cirl^Rescue^, Cirl^7TM^* and *Cirl^1TM^* larvae were recorded in free locomotion. The average midpoint speed of each genotype in 5 s bins and the 95% confidence interval of the recording is displayed. *Cirl^KO^* animals were slower than the *Cirl^Rescue^* ^22^. This phenotype was rescued to and above control in *Cirl^7TM^* and *Cirl^1TM^* animals, respectively. **(B)** Quantification of average midpoint speed (normalized to *Cirl^Rescue^*) of each animal cohort over 2 min recordings. Related to Table S3. **(C)** Schematic of action current recording configuration. **(D, E)** Experimental protocol. Pull lengths (0.03, 0.1, 0.3, 1, and 3 μm) were applied for 500 ms. Differential reactions of the lch5 neurons as an increase of AC frequency after applying and releasing the pull are shown schematically (D) and in representative traces (E). ACs within the first 50 ms (gray box) after stimulation onset were quantified to calculate the AC frequency. F = Force. **(F-H)** Quantification of AC frequencies from mechano-stimulated lch5 neurons. Upper panels: Sample AC traces in response to a 30-nm pull length. The gray shades indicate the 50 ms time period from which the AC frequency was quantified. Lower panels: Quantification of lch5 neuron activity at different pull lengths (in μm: 0.03, 0.1, 0.3, 1, 3). Data are displayed as mean ± SEM. P-values denote statistical difference between *Cirl^Rescue^* and *Cirl^7TM^* (F), *Cirl^1TM^* (G) and *Cirl^7TM+1TM^* (H) at 30 nm pull length. Color code: *Cirl^Rescue^* white, *Cirl^KO^* gray, *Cirl^7TM^* blue, *Cirl^1TM^* green, *Cirl^7TM+1TM^* orange. Related to Figure S4 and Tables S4,5.

The lch5 neurons of *Cirl^Rescue^* larvae responded to a 0.03 μm pull with a mean AC frequency of 140 Hz. The frequency increased with rising pull lengths to 325 Hz at a 3 μm pull (Δ_0.03-3μm_ = 185 Hz; Figure 3F-H, Table S4), which shows that lch5 neurons can differentiate between these two pull lengths, i.e., stimulation strengths. In contrast, lch5 neurons of *Cirl^KO^* were much less capable of discriminating between different pull lengths (Δ_0.03-3μm_ = 26 Hz; Figure 3E, Table S4). Hence, in *Cirl^KO^* the lch5 output signal relayed to the central nervous system remains virtually unaltered although the actual stimulation intensity has drastically changed. This finding is consistent with the discrimination deficits of *Cirl^KO^* between different vibration frequencies in *Cirl^KO^* ^22^. Strikingly, lch5 responses from *Cirl^7TM^* or *Cirl^1TM^* neurons were similar to those of *Cirl^KO^* animals in that each exhibited a discrimination deficit (Figure 3F,G, Table S4,5). Importantly, appending a biochemical tag at Cirl^1TM^ C-terminus (Figure S4, Table S5) or Cirl^7TM^ third intracellular loop had no impact^21^.

To test for a co-requirement and to rule out technical issues for the observed mechanosensation deficits in *Cirl^7TM^* and *Cirl^1TM^* mutants, we generated trans-heterozygous *Cirl^1TM^/Cirl ^7TM^* (*Cirl^1TM+7TM^*) animals reconstituting the natural Cirl isoform repertoire. Reassuringly, lch5 responses in these animals (Δ_0.03-3μm_ = 106 Hz; Figure 3H, Tables S4,5) were comparable to *Cirl^Rescue^* despite containing only a single allele copy of each splice species. Collectively, this shows that Cirl^7TM^ (GPCR-like) isoforms require Cirl^1TM^ (non-GPCR) isoforms to tell different mechanical input intensities apart.

### Sound stimulation evokes activity in select lch5 neurons that necessitates Cirl^7TM^ and Cirl^1TM^ proteins

Cirl is involved in lch5-dependent sound detection^22^. With the tonotopic organization of the mammalian auditory system in mind, we asked if individual lch5 neurons respond differently toward the same sound stimulus and whether different Cirl isoforms are relevant. Therefore, we performed Ca^2+^ imaging as a proxy for neuronal lch5 activity employing the genetically encoded Ca^2+^ indicator *UAS-jGaMP7*^36^ expressed under *iav-GAL4* control^37^ (Figure 4A,B). Interestingly, in *Cirl^Rescue^* 90 dB sound pressure level (SPL) stimuli repeatedly elicited the largest response in the most ventral neuron (= neuron 1; Figure 4C, Table S6), implying that lch5 neurons are differentially tuned. Hence, to investigate Cirl isoform function on single-cell level, we quantified sound-induced Ca^2+^ signals in neuron 1. Consistent with the elevation in AC frequency measured electrophysiologically (Figure 3F-H), neuron 1 of *Cirl^KO^* showed markedly increased Ca^2+^ signals when challenged with sound (Figure 4D-F, Table S7). Importantly, loss of either isoform species had similar effects corroborating the physiological necessity of the non-GPCR Cirl^1TM^ proteins (Figure 4D-E). Note that ΔF/F_0_ values from *Cirl^7TM^*^+*1TM*^ animals (containing only a single gene copy of *Cirl^7TM^* and *Cirl^1TM^* allele, respectively) ranged between those of *Cirl^KO^* and *Cirl^Rescue^* (Figure 4F) suggesting that the amount of Cirl protein is a limiting factor for the acoustic sensitivity of neuron 1. Ca^2+^ signals in neuron 2 and 4 seemed indistinguishable between genotypes (Figure 4G,H, Table S8). In sum, spatial analyses of mechano-dependent Ca^2+^ signals uncovered differential activity profiles of lch5 neurons and further underpins that Cirl function hinges on the presence of the Cirl^1TM^ non-GPCR isoforms.

**Figure 4.**
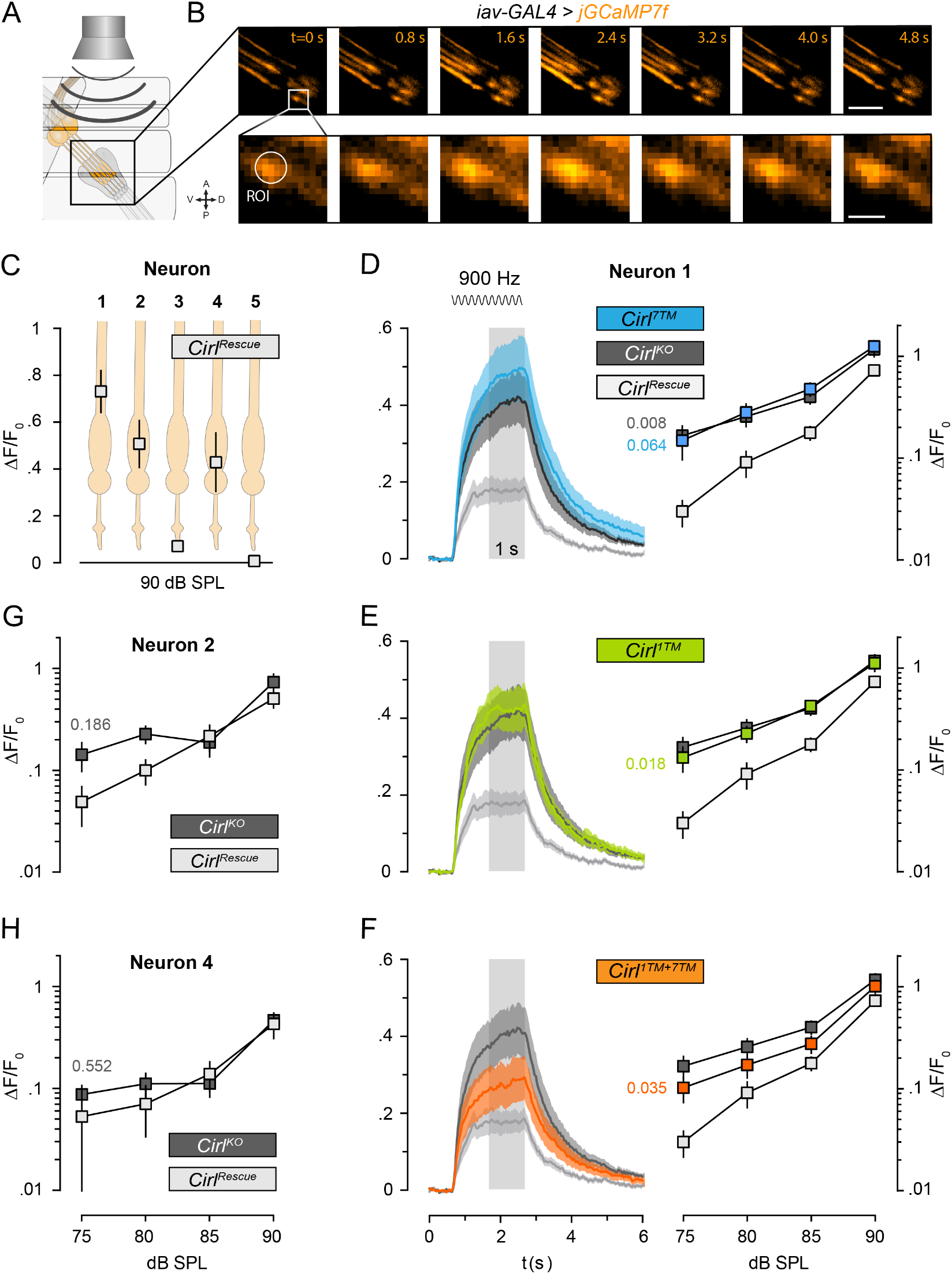
Sound stimulation evokes activity in select lch5 neurons that necessitates Cirl^7TM^ and Cirl^1TM^ proteins. **(A)** Schematic of Ca^2+^ imaging configuration. Ca^2+^ indicator jGCaMP7f (orange) outlines lch5 neurons. **(B)** Representative images of a time series (t = 0 – 4.8 s) of Ca^2+^ signals visualized from larval lch5 organs (*iav-GAL4 > jGaMP7f*). A 900 Hz, 85 dB SPL sound was applied for 2 s (from 0.7 s to 2.7 s). To quantify Ca^2+^ signals, fluorescence intensities within a circular area with a 5-pixel diameter (~ 815 nm) were measured in the distal part of the dendrite. Region of interest (ROI) in neuron 1 shown exemplarily. Scale bars: 5 μm (upper), 1 μm (lower). **(C)** Schematic of lch5 neurons (orange) overlaid by Ca^2+^ signal average ΔF/F_0_ (square) of the plateau phase (1 s indicated by gray shade in (D)) measured from *Cirl^Rescue^*, shown as mean ± SEM. Neuron 1 is the most ventral neuron. Stimulation intensity: 90 dB SPL. Related to Table S6. **(D-F)** Quantification of Ca^2+^ signals measured within the dendrite of neuron 1. Left panels: Mean time course of ΔF/F_0_ ± SEM (each genotype n = 10). Stimulation period (2 s, 900 Hz, 85 dB SPL) is indicated by a sine wave. Right panel: To quantitatively compare sound evoked Ca^2+^ signals between genotypes, ΔF/F_0_ averages and SEMs were calculated from the plateau phase (1 s) at different stimulation intensities (75, 80, 85, 90 dB SPL). P-values shown are derived from statistical comparisons at 75 dB SPL between *Cirl^Rescue^* and the three mutant genotypes (n=10). Color code: *Cirl^Rescue^* white, *Cirl^KO^* gray, *Cirl^7TM^* blue, *Cirl^1TM^* green,*Cirl^7TM/1TM^* orange. Related to Table S7. **(G, H)** Quantification of the Ca^2+^ signals in neurons 2 and 4. P-values from comparison of *Cirl^Rescue^* and *Cirl^KO^*. In (G), at 75 dB SPL, n = 4 and n = 7 for *Cirl^Rescue^* and *Cirl^KO^*, respectively. In (H), at 75 dB SPL, n = 5 for both *Cirl^Rescu^*^e^ and *Cirl^KO^*, respectively. Related to Table S8.

### Cirl^1TM^ and Cirl^7TM^ form a heteromeric complex

The previous results collectively suggest that Cirl^1TM^ and Cirl^7TM^ are co-required to modulate the neuro-mechanical response of lch5 neurons. Consequently, we asked whether Cirl^1TM^ and Cirl^7TM^ functions merely intersect or if Cirl isoforms assemble signaling complexes to shape mechanosensation. To test for a direct Cirl^1TM^-Cirl^7TM^ interaction, we used HEK293T cells as a heterologous expression system. First, we conducted ELISA to quantify the expression levels of specific Cirl isoforms using an N-terminal hemagglutinin (HA) tag^21,38^ and found Cirl^1TM^ isoform (-F/K) more abundantly at the cell surface than Cirl^7TM^ isoforms (-B/H and -E) (Figure 5A). GPS-dependent autoproteolytic cleavage is relevant for trafficking of several aGPCR (Figure 5B)^39–41^. Therefore, we wondered whether self-cleavage affects cell surface expression of different Cirl isoforms and, if so, whether the position of the cleavage-disabling mutation (H>A^GPS-2^ or T>A^GPS+1^) is relevant. ELISA showed that cleavage-deficiency increased total and surface expression of Cirl^7TM^-E receptors (Figure 5C). In contrast, the cell surface pool of Cirl^1TM^-F/K isoforms remained unaltered irrespective of autoproteolytic cleavage competence (Figure 5C). Collectively, this shows that the faculty for self-cleavage and the layout of the CTF impacts cell surface trafficking of Cirl isoforms.

**Figure 5.**
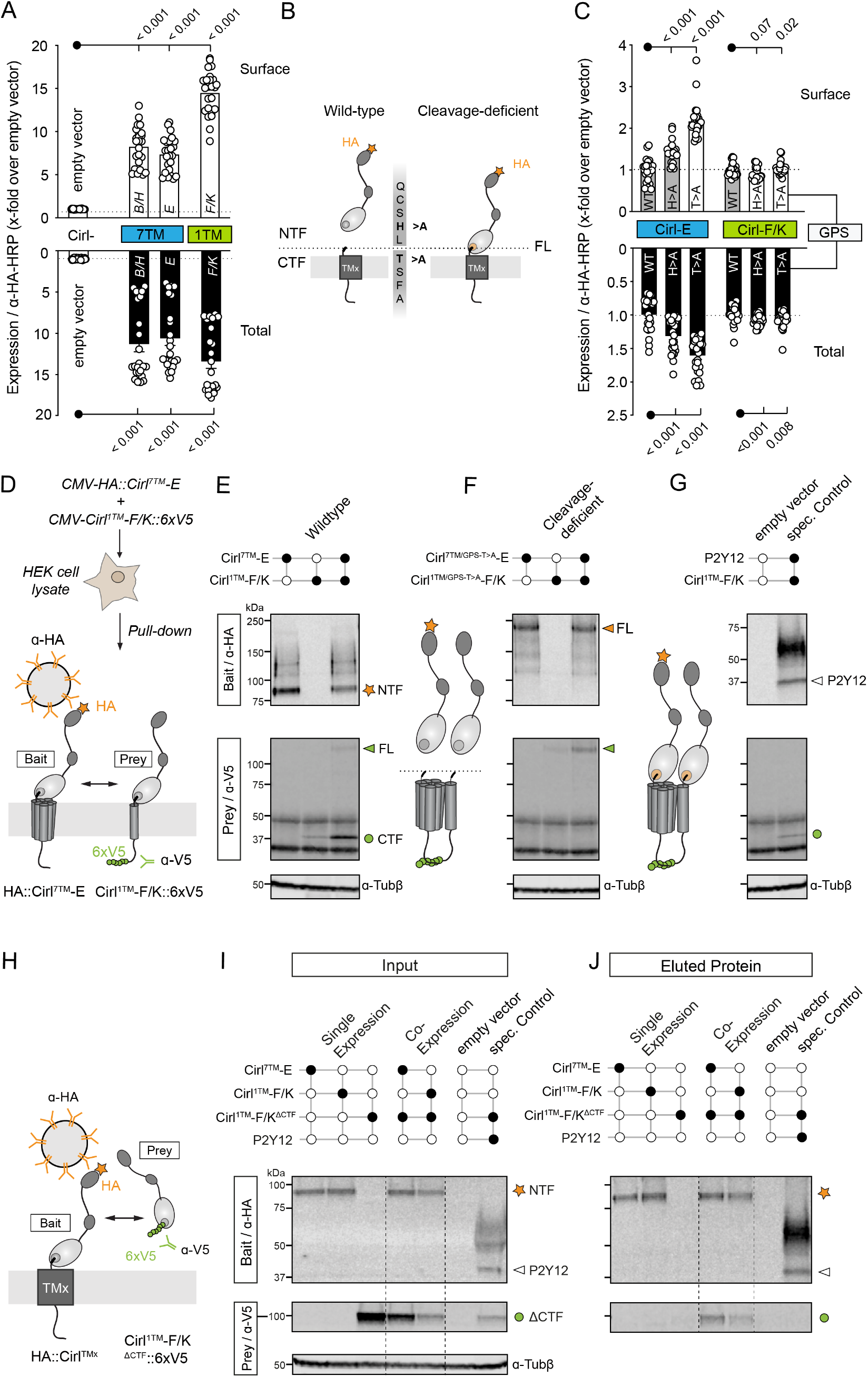
Cirl^1TM^ and Cirl^7TM^ form a heteromeric complex. **(A)** Quantification of total and cell surface expression of individual Cirl isoforms by ELISA. Cirl detection was done using a N-terminal HA tag (HA∷Cirl-X). Data are shown as mean ± SEM. Related to Table S9. **(B)** Schematic of HA-tagged wildtype and cleavage-disabled Cirl. **(C)** ELISA of wildtype and cleavage-deficient Cirl-E and -F/K. Data of cleavage-deficient Cirl variants were normalized to respective wildtype receptor and are shown as mean ± SEM. Related to Table S9. **(D)** Schematic of pull-down experiments shown in E-G. **(E-G)** Immunoblot of a pull-down experiment using wildtype (E) and cleavage-deficient (F) Cirl. HA-tagged Cirl^7TM^ receptor (HA∷Cirl-E) served as bait and was immobilized/detected using α-HA antibody (upper row); Cirl^1TM^ prey (Cirl^1TM^-F/K∷V5) was detected using α-V5 antibody (lower row). FL and NTF of Cirl^7TM^-E are indicated by orange arrowhead and orange star, respectively. FL and CTF of Cirl^1TM^-F/K are indicated by green arrowhead and green circle, respectively. All samples were run side-by-side on the same gel. (G) Empty vector: beads incubated with lysate from empty vector transfected cells. Specificity control: P2Y12 receptor (white arrowhead) + Cirl^1TM^-F/K. Unspecific binding of Cirl^1TM^-F/K∷V5 to the beads returned background signal (green circle), which was below that of co-precipitated Cirl^1TM^-F/K (E). Tubulin served as loading control (α-Tub*β*). **(H)** Schematic of pull-down experiments shown in I and J. HA-tagged Cirl^TMx^ protein (HA∷Cirl^7TM^-E or HA∷Cirl^1TM^-F/K) served as bait for soluble Cirl^1TM^-NTF(Cirl^1TM^-F/K^ΔCTF^∷6xV5). **(I, J)** Immunoblots from pull-down experiments show that isoform assembly involves the NTF of Cirl. Input samples (I) and eluted protein (J). Cirl^TMx^ was immobilized/detected using α-HA antibody (upper row, orange star). Detection of Cirl^1TM^-F/K^ΔCTF^ (green circle) was done using α-V5 antibody (lower row). Empty vector: beads incubated with lysate from empty vector transfected cells. Specificity control: P2Y12 receptor (white arrowhead) + Cirl^1TM^-F/K^ΔCTF^. Tubulin served as loading control (α-Tub*β*).

To test whether Cirl^7TM^-Cirl^1TM^ complexes assemble, we overexpressed HA∷Cirl^7TM^-E with and without C-terminally tagged Cirl^1TM^-F/K proteins (Cirl^1TM^-F/K∷V5) under *CMV* promoter (*CMVp*) and performed pull-downs. HA∷Cirl^7TM^-E served as bait to capture Cirl^1TM^-F/K∷6xV5 protein (Figure 5D). Western blot analyses of cell lysates from pull-down experiments showed Cirl^7TM^-E-specific enrichment of Cirl^1TM^-F/K-CTF (~ 38 kDa; Figure 5E) lending support for a direct interaction of Cirl^7TM^ and Cirl^1TM^ isoforms (input samples shown in Figure S4B-D). We repeated the experiment employing cleavage-deficient (T>A^GPS+1^) isoform versions and were able to enrich FL Cirl^1TM/GPS-T>A^-F/K (~ 117 kDa; Figure 5F). We detected faint background signals when Cirl^1TM^-F/K∷V5 was overexpressed alone or together with the class A GPCR P2Y12. However, the intensities of these signals were well below those of Cirl^1TM^-F/K when enriched by Cirl^7TM^-E (Figure 5E,G). Importantly, Cirl^7TM^-E could also be captured in an inverse experimental set-up using Cirl^1TM^-F/K as bait (Figure S4H-J).

Interestingly, when we placed *Cirl^1TM^* under transcriptional control of a single *UAS*-promotor (Figure S4E) and induced Gal4 expression using the *actin5C*-promoter^42^, we detected Cirl^1TM^-F/K in input samples only when Cirl^7TM^-E was co-expressed (Figure S4F). Importantly, in this modified pull-down set-up Cirl^1TM^ also co-immunoprecipitated with Cirl^7TM^ cleavage-independently (Figure S4G). Note that we used ADGRE5/CD97 here as mock bait instead of P2Y12. This was done as an additional measure to ensure specificity of Cirl complex formation (Figure S4G).

Finally, to gain a first insight into Cirl isoform interaction interface(s), we expressed a soluble NTF-only version of Cirl^1TM^-F/K *i.e*. lacking the CTF (Cirl^1TM^-F/K^ΔCTF^, Figure 5H). Cirl^1TM^-F/K^ΔCTF^ also formed a complex with Cirl^7TM^-E (Figure 5I,J), which suggests that Cirl^1TM^-NTF is part of the binding interface between different Cirl isoforms. Reassuringly, Cirl^1TM^-F/K^ΔCTF^ engaged in a homomeric interaction with membrane anchored Cirl^1TM^-F/K, which was expected as it shares NTF layout with Cirl^7TM^-E (Figure 5J). Collectively, these findings render homomeric interaction between Cirl isoforms possible and show that Cirl^1TM^ molecules can physically interact with Cirl^7TM^ receptor in an NTF-dependent and cleavage-independent fashion.

### Cirl signals through the Gα_o_ subunit to attenuate cAMP quantity

Previous work showed that Cirl affects lch5 mechanosensing through a cAMP-dependent signaling route^21^. Cirl^1TM^ proteins are ill-equipped to couple G proteins and thus are most likely unfit to act through inhibition of cAMP formation. This in combination with the isoform interaction led us to interrogate if aberrant lch5 mechanosensing in *Cirl^1TM^* mutants can be reversed through pharmacological inhibition of adenylyl cyclases (ACyc), enzymes responsible for G protein-dependent cAMP production (Figure 6A). Similar to *Cirl^KO^* and *Cirl^7TM^* lch5 neurons, application of 100 μM SQ22536, an ACyc inhibitor, indeed rescued mechanically-evoked AC frequencies in *Cirl^1TM^* (Figures 6B-D, S5A, Table S10,11) suggesting a role of Cirl^1TM^-Cirl^7TM^ complexes in the suppression of cAMP levels.

**Figure 6.**
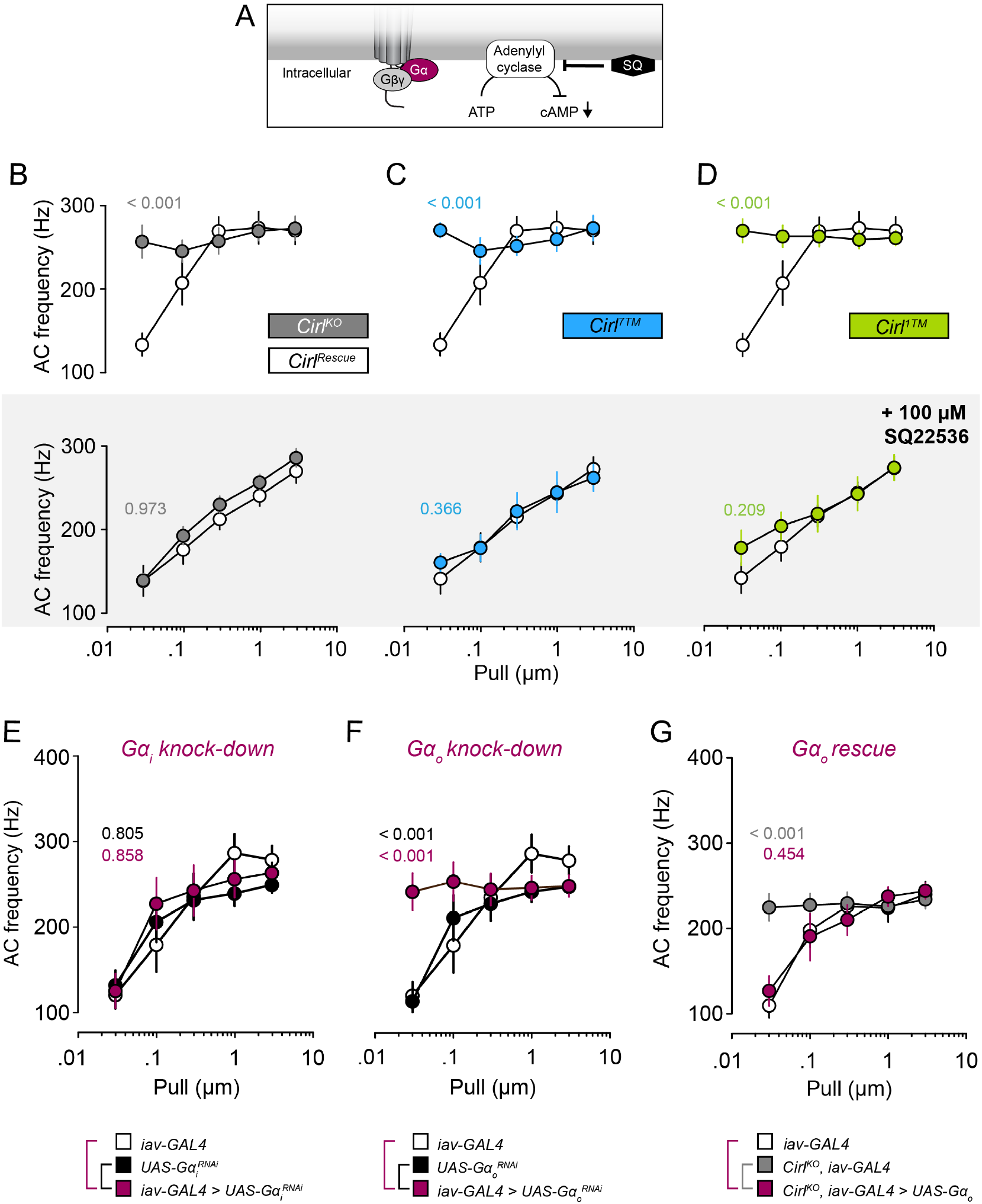
Cirl signals through Gα_o_ subunit to attenuate cAMP quantity. **(A)** Schematic of pharmacological inhibition of adenylyl cyclases (ACyc) by SQ22536, which decreases cAMP levels. **(B-D)** Quantification of AC frequencies from mechanically stimulated lch5 neurons before and after treatment with ACyc inhibitor. (B-D) Show the response of *Cirl^KO^* (B), *Cirl^7TM^* (C) and *Cirl^1TM^* (D) compared to *Cirl^Rescue^* before treatment with the ACyc blocker SQ22536. Gray-shaded area shows lch5 responses of the three mutants 10 min after addition of 100 μM SQ22536. P-values denote statistical differences of AC frequencies at a pull length of 30 nm between *Cirl^Rescue^* and the three mutants. *Cirl^Rescue^*, white; *Cirl^KO^*, gray; *Cirl^7TM^*, blue; *Cirl^1TM^*, green. Related to Figure S5A and Table S10. **(E,F)** Quantification of AC frequencies from mechanically stimulated *Gα_i_* or *Gα_o_*-deficient lch5 neurons (*iav-GAL4* > *UAS-Gα_i_^RNAi^ or UAS-Gα_o_^RNAi^*). (E) *Gα_i_* knock-down leaves AC frequencies unaffected. (F) *Gα_o_* knock-down leads to an increase in AC frequencies, phenocopying the *Cirl^KO^*. P-values are derived from the statistical comparison of the AC frequencies at 30 nm pulls between *iav-GAL4* > *UAS-Gα_i_^RNAi^* or *iav-GAL4 > UAS-Gα_o_^RNAi^* (in magenta) and *iav-GAL4* (in white) or *UAS-RNAi* controls (in black). Related to Figure S5B and Tables S12,13. **(G)** Quantification of AC frequencies from mechanically stimulated lch5 neurons overexpressing Gα_o_ in the absence of *Cirl*. Lch5-specific Gα_o_ expression rescues elevated AC frequency of *Cirl^KO^* back to control levels. P-values are derived from the statistical comparison of the AC frequencies at 30-nm pulls between *Cirl^KO^*, *iav-GAL4* > *UAS-Gα_o_*, (in magenta) and *iav-GAL4* (in white) or *Cirl^KO^*, *iav-GAL4* (in gray). Related to Figure S5C and Table S16. **(B-G)** Data are shown as mean ± SEM (each genotype n = 10).

Next, we asked which Gα protein species acts downstream of Cirl to suppress neuronal cAMP levels. Gα_i_ protein is expressed in developing lch5^43^, inhibits ACycs and thus reduces cAMP^44^. However, when we knocked-down the *Gα_i_* gene in lch5 using two different RNAi lines (*iav-GAL4 > UAS-Gα_i_-RNAi-1 and −2*), we found no effect on lch5 mechanoresponses (Figures 6E, S5B, Table S12,13,14). Despite the controversial view on how Gα_o_ modulates cAMP levels, this subunit is known to play vital roles in different neuronal contexts^45–49^. We measured the mechano-response of *Gα_o_*-depleted lch5s (*iav-GAL4 > UAS-Gα_o_-RNAi*), which phenocopied the deficit of null and isoform-specific *Cirl* mutants (Figure 6F, Table S12,13). Affirmingly, lch5-specific neuronal overexpression of Gα_o_ in lch5 neurons of *Cirl^KO^* animals (*Cirl^KO^, iav-GAL4 > UAS-Gα_o_*)^48^ rescued the AC frequency (Figure 6G, Table S15,16). Together these data show that Cirl acts through Gα_o_ but not Gα_i_ and suggest that 7TM-isoforms cooperate with 1TM protein, presumably to decrease neuronal cAMP levels.

### Mammalian 1TM and 7TM GPR126/G6 isoforms interact

Finally, we tested if isoform-specific complex assembly is a common phenomenon also observed for mammalian aGPCR. GPR126/G6 is an aGPCR with abundant expression in the nervous system, heart and bone^50^, whose splice spectrum includes 1TM and 7TM-containing transcript variants^27^. For co-immunoprecipitation experiments, we appended the N terminus of G6^7TM^ with an HA tag (HA∷G6^7TM^) and the C terminus of G6^1TM^ with V5 tags (G6^1TM^∷V5, Figure 7A). Both proteins were robustly expressed in HEK293T cells (Figure 7B,C) and an extracellular domain fragment (ECD^fragment^) of HA∷G6^7TM^ and an FL G6^1TM^∷V5 protein were found in the input sample (Figure 7B). Intriguingly, a FL and an intracellular fragment (ICD^fragment^) of G6^1TM^∷V5 was immobilized by HA∷G6^7TM^ bait when co-expressed. When GAIN domain cleavage of HA∷G6^7TM^ and G6^1TM^∷V5 was disabled by mutation, co-immunoprecipitation still captured FL G6^1TM^∷V5 protein but, as expected, no ICD^fragment^ (Figure 7C). Importantly, G6^1TM^∷V5 using the ADP receptor P2Y12 receptor as bait was not detected confirming specificity of the G6^1TM^-G6^7TM^ interaction (Figure 7C).

**Figure 7.**
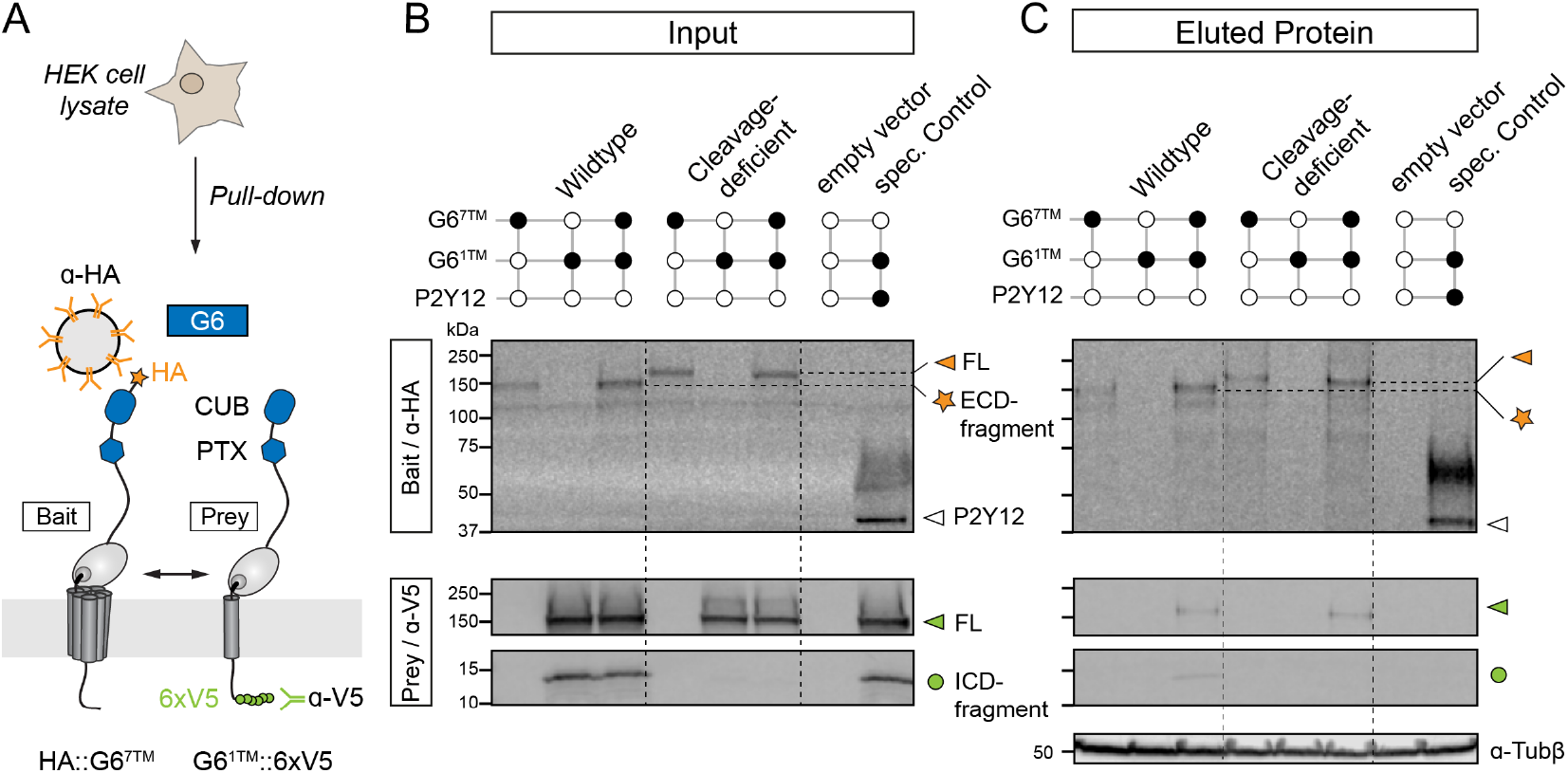
Mammalian 1TM and 7TM GPR126/G6 isoforms interact. **(A)** Schematic of the experimental configuration. N-terminally tagged HA∷G6^7TM^ immobilized via α-HA functionalized magnetic beads served as bait to capture G6^1TM^∷V5. **(B, C)** Immunoblots of G6 pull-down experiment. Input (B); eluted protein (C). Upper row (bait): α-HA antibody recognizes FL (orange arrowhead) and an ECD fragment of HA∷G6^7TM^ (orange star). Lower row (prey): α-V5 antibody recognizes FL (green arrowhead) and an ICD fragment of G6^1TM^∷V5 (green circle). G6 isoforms were expressed under *CMVp*. HA-tagged P2Y12 was used as control bait to ensure specificity of the interaction between G6 isoform. Tubulin served as loading control (α-Tub*β*).

Collectively, our findings suggest that the formation of isoform complexes may be a general phenomenon shaping aGPCR biology in different species.

## DISCUSSION

While mechanosensing through ionotropic mechanosensors is well recognized^16^, it is largely unknown how metabotropic mechanosensors establish their sensitivity. Here, we show that *Cirl* uses intron retention as a physiological strategy to produce non-GPCR-coding Cirl^1TM^ isoforms. These form protein complexes with GPCR-like Cirl^7TM^ receptors to tune the mechanosensitive abilities of sensory neurons (Figures 3–5). At first glance, this is reminiscent of receptor modifying proteins (RAMP)^51^, which however, are encoded in genes separate from those GPCR that they modify.

Intron retention is a major alternative splicing mode^52^, which is poorly understood^53^ and even often regarded a consequence of erroneous splicing counteracted by nonsense-mediated mRNA decay^54^. Contrarily, intron retention was reported to control developmentally regulated genes^55^ and to curtail the abundance of protein-coding transcripts^56^. Importantly, intron retention rarely has been seen through to the level of predicted protein isoforms encoded in mature transcripts with retained introns^57^. Alternative splicing of *Cirl* generates *Cirl^7TM^* transcripts (removal of intron 6) and *Cirl^1TM^* transcripts (retention of intron 6) that encode an alternative intracellular tail and a premature stop codon (Figures 1 and S1). We found both transcript species co-expressed throughout development (Figure 1). In sensory lch5 neurons of *Drosophila* larvae, Cirl^7TM^ and Cirl^1TM^ isoforms are located in close proximity to the ciliary mechanosensing apparatus and are equally required for the detection of different mechanical stimulus qualities: sound and stretch at different intensities (Figures 2–4). Interestingly, exclusive Cirl^7TM^ or Cirl^1TM^ pools exist in some subcellular regions implying functions outside of the Cirl^1TM^-Cirl^7TM^ complex (Figure 2). The finding that Cirl^7TM^ suffices to restore locomotion speed to control values lends support to this notion (Figure 3). The functional relevance of different isoforms may explain why previous rescue attempts of aGPCR knockout animal models with transgenes that only encoded GPCR-like 7TM proteins were futile, while rescue strategies with genomic transgenes (retaining the ability to produce GPCR-like and non-GPCR isoforms) were successful^22,58,59^.

Dimer and oligomer formation between canonical 7TM-GPCR has been controversially discussed for many years^60–63^. While dimerization of GPCR such as GABAB and mGlu receptors occurs between 7TM protomers and is required for membrane localization, ligand binding and signal transduction^64,65^, we show in this study that complex formation manifests between 7TM and 1TM protomers of the aGPCR Cirl from *Drosophila* and ADGRG6 from mouse (Figures 5 and 7). This finding is in agreement with a previous intermolecular complementation experiment in which co-expression of a wildtype LAT-1 CTF (7TM protein) and a chimeric protein containing the ECD of the *C. elegans* latrophilin homolog LAT-1 fused to the transmembrane unit of PAT-3 (1TM protein) was sufficient to restore full receptor activity. Moreover, the LAT-1 ECD was found to form stable homodimers^66^, which is in line with our finding that Cirl isoforms complex via their NTFs into heteromeric and potentially homomeric assemblies (Figure 5).

Recent structural evidence shows direct engagement of the tethered agonist (TA)/Stachel region with the 7TM domain for several aGPCR^67–70^. The newfound functionality of Cirl 7TM-1TM dimers implies that the GAIN domain of at least the 1TM protomer can exist in a steric configuration that does not require TA-7TM-interaction. Alternatively, TA-7TM engagement may occur between rather than within protomers as implied before^66^.

We show that Cirl signals through Gα_o_, but not Gα_i_ proteins to reduce cAMP levels and to adjust mechano-evoked AC frequencies of sensory lch5 neurons (Figures 6 and S5). Brain-derived rat Latrophilin-1 co-precipitates with Gα_o_ protein^71^, the most abundant yet puzzling Gα subunit in nervous tissue^45,72^. Whether Gα_o_ protein couples to Cirl^7TM^ or the Cirl^1TM^-Cirl^7TM^ complex is unclear, but the fact that a loss of either of the three proteins results in a similar phenotype indicates the necessity of Cirl^1TM^ to efficiently couple Gα_o_ to Cirl^7TM^ (Figure 6). Alternatively, Cirl^1TM^ may act through another Gα_o_-coupled receptor in lch5, however, due to absence of additive effects in *Cirl^KO^*, i.e. loss of both *Cirl^7TM^* and *Cirl^1TM^*, this scenario is less likely. Whether Gα_o_ impacts cAMP concentrations directly is currently debated^49^, but it is tempting to speculate that Gα_o_ as a potential liberator of Gβγ subunits affects ion channel gating and indirectly cAMP levels^73^. This model fits previous work, where modulation of mechanogating properties of transient potential receptors through Cirl was shown^21^.

Ca^2+^ imaging of acoustically stimulated lch5 neurons showed robust differences in neuronal activity across the five lch5 cells (Figure 4). Upon acoustic stimulation the highest Ca^2+^ signals were recorded from neuron 1, medium signals in neurons 2 and 4, while neurons 3 and 5 were unresponsive. This could stem from morphological variability of lch5 neurons or differing sensitivity thresholds across cells, ultimately contributing to the mechanical responsiveness of the neurons. While Ca^2+^ response of neuron 1 relied on the presence of both Cirl^1TM^ and Cirl^7TM^, it is unclear if and how Cirl or which isoforms are involved in weaker signals recorded from neurons 2 and 4.

Alternative splicing is particularly prominent in nervous tissues^74,75^ and may be causally related to its tremendous information processing power and plasticity. Here, we present proof that alternative splicing of aGPCR is a physiological strategy that establishes sensitivity towards defined mechanical stimuli in individual neurons. This discovery is underpinned by the utility of aGPCR complex formation through the expression and unexpected signaling role of non-GPCR isoforms. This offers an explanation for the versatile splice repertoire of aGPCR genes as a source of modulators for metabotropic signals transduced by aGPCR proteins. Such interactions may prove useful as means to interfere with aGPCR signaling and mechanosensitivity in the nervous system and other organs, which may pave the way to novel pharmacological strategies.

## EXPERIMENTAL PROCEDURES

### *Drosophila* culture and stocks

*Drosophila* were cultured on standard cornmeal food at 25 °C. The following strains were generated in this study:

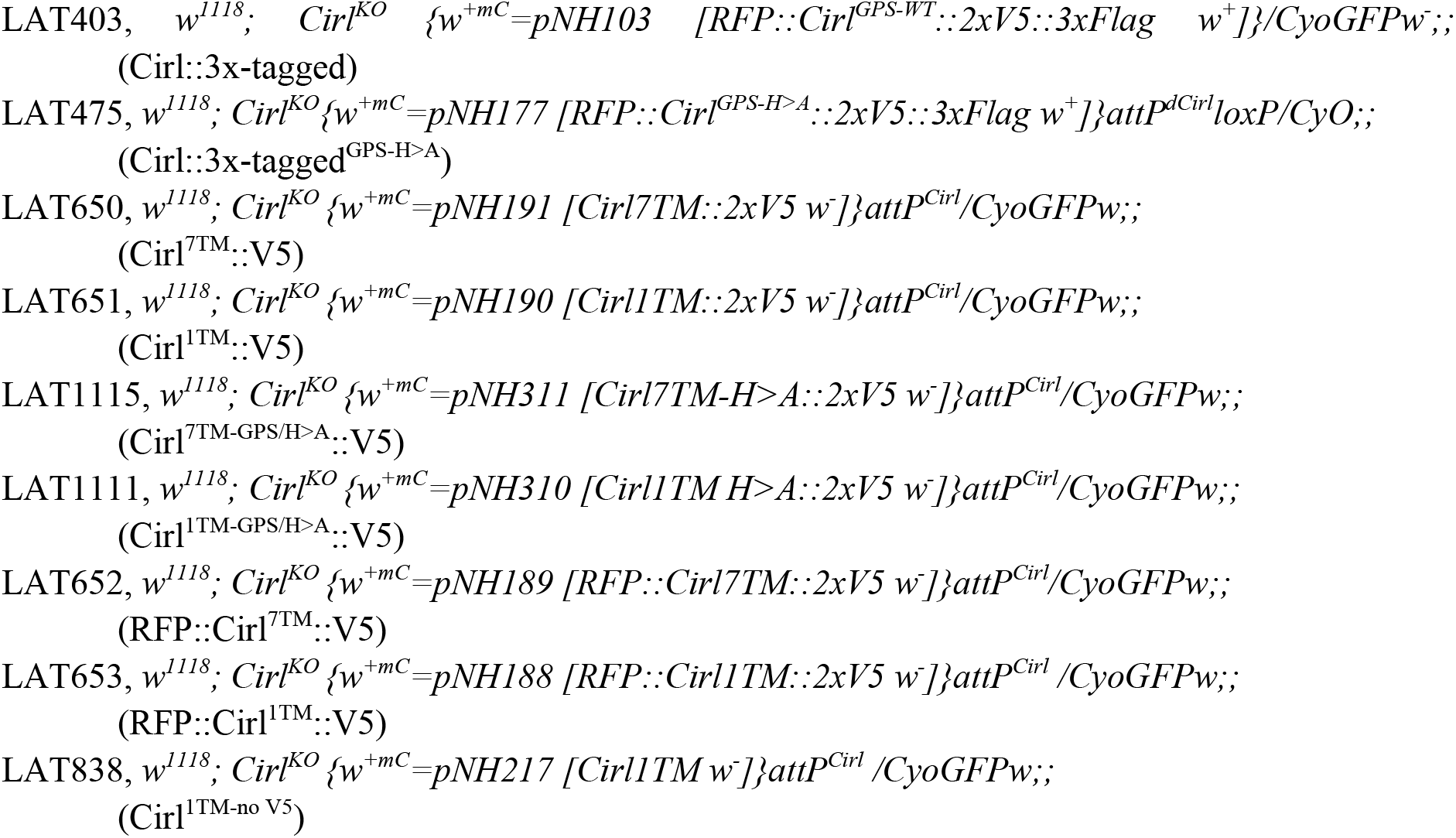

The following strains were previously generated or generated elsewhere:

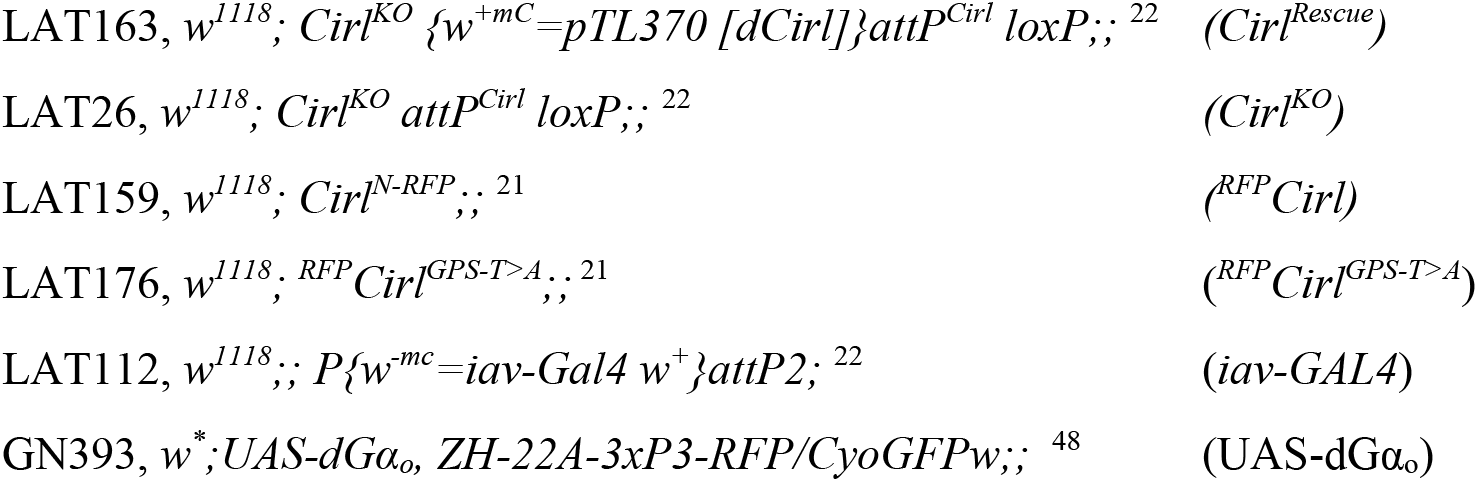

Bloomington Stock Center (NIH P40OD018537) or Vienna *Drosophila* Research Center ^76^.

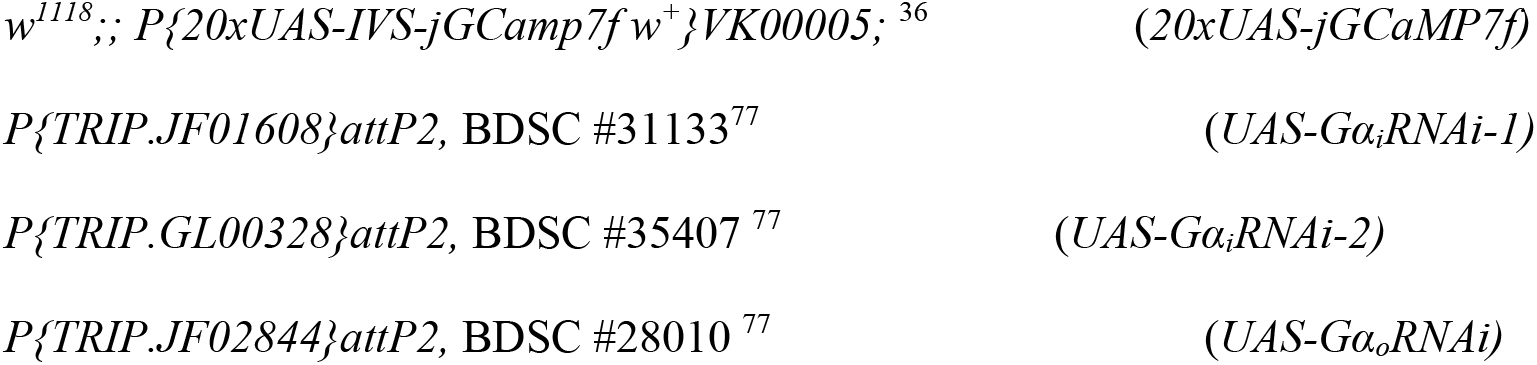

### Molecular cloning

*Cirl* cDNAs were N-terminally complemented with a mammalian Kozak sequence (gccacc), followed by the start codon, signal peptide (SP) of mammalian ADGRG1/GPR56, an HA tag (*HA∷Cirl-X)* and are contained in the *CMV* promoter-containing pcDNA3.1 backbone (Table S1). Cirl^1TM^ constructs used for immunoprecipitation experiments lacked HA tag and contained six V5 tags at their C termini (Cirl^1TM^∷V5). Primers used for cloning of *Cirl* and *ADGRG6/GPR126* cDNA constructs are listed in Table S2. Plasmid amplification was done via transformation in *E. coli* (XL1-Blue, DH5α or EPI400) and DNA isolation via Midiprep (NucleoBond® Xtra Midi, Macherey & Nagel). The Ranger (Bioline, #BIO21121) or Q5 (New England Biolabs, #M0491S) high-fidelity proof-reading DNA polymerases were utilized for all PCR-based cloning steps. Initial insert verification was done by restriction analyses. To ensure absence of errors each PCR-amplified region was sequenced completely.

Construction of *Drosophila* transgenes was done as follows:

#### pNH103 (Cirl∷3x-tagged)

Primers *nh_199F/259R* were used to amplify a 4.9-kbp fragment from *pMN21* (genomic 1.9 kbp *Cirl* subclone) and to introduce *Sgf*I and *Stu*I restriction sites at the 3’ and 5’ end, respectively.

2xV5 tag was generated using *nh_260F/261R*, which contained a *Sgf*I and *Stu*I sites for subsequent ligation with 4.9-kbp fragment (resultant subclone *pNH101*). *Cirl∷2xV5* fragment was transplanted from *pNH101* into *pMN24* (genomic *RFP∷Cirl∷3xFlag* construct) via *Pac*I and *Spe*I restriction sites resulting in *pNH103*.

#### pNH177 (Cirl∷3x-tagged^GPS/H>A^)

Primers *mn_38F/39R* were used to substitute the histidine codon of the canonical GPS sequence for an alanine codon through site-directed mutagenesis of *pNH101*, which resulted in *pNH176*. *Pac*I/*Spe*I fragment of *pNH176* was cloned into *pNH103* resulting in *pNH177*.

#### pNH188 (RFP∷Cirl^1TM^∷2xV5)

Genomic sequence encoding *RFP∷Cirl^1TM^∷2xV5* (~ 3.9 kbp) in pMK backbone was synthesized by ThermoFisher/GeneArt (resulting construct *pNH186*). A 3.9-kbp fragment was cloned into the expression vector using *Mlu*I and *BstB*I sites to generate *pNH188*.

#### pNH189 (RFP∷Cirl^7TM^∷2xV5)

Genomic sequence encoding *RFP∷Cirl^7TM^∷2xV5* (~ 5.7 kbp) in pMK backbone was synthesized by ThermoFisher/GeneArt (resulting construct *pNH187*). A 5.7-kbp fragment was cloned into expression vector using *Mlu*I and *Aat*II sites to generate *pNH189*.

#### pNH190 (Cirl^1TM^∷2xV5)

*Age*I-based removal of mRFP sequence and subsequent re-circularization of *pNH188* resulted in *pNH190*.

#### pNH191 (Cirl^7TM^∷2xV5)

*Age*I-based removal of mRFP sequence and subsequent re-circularization of *pNH189* resulted in *pNH191*.

#### pNH217(Cirl^1TM^ no tags)

Engineered by GenScript.

#### pNH310 (Cirl^1TM-GPS/H>A^∷2xV5)

Engineered by GenScript.

#### pNH311 (Cirl^7TM-GPS/H>A^∷2xV5)

Engineered by GenScript.

#### Primers (5’-3’ orientation)

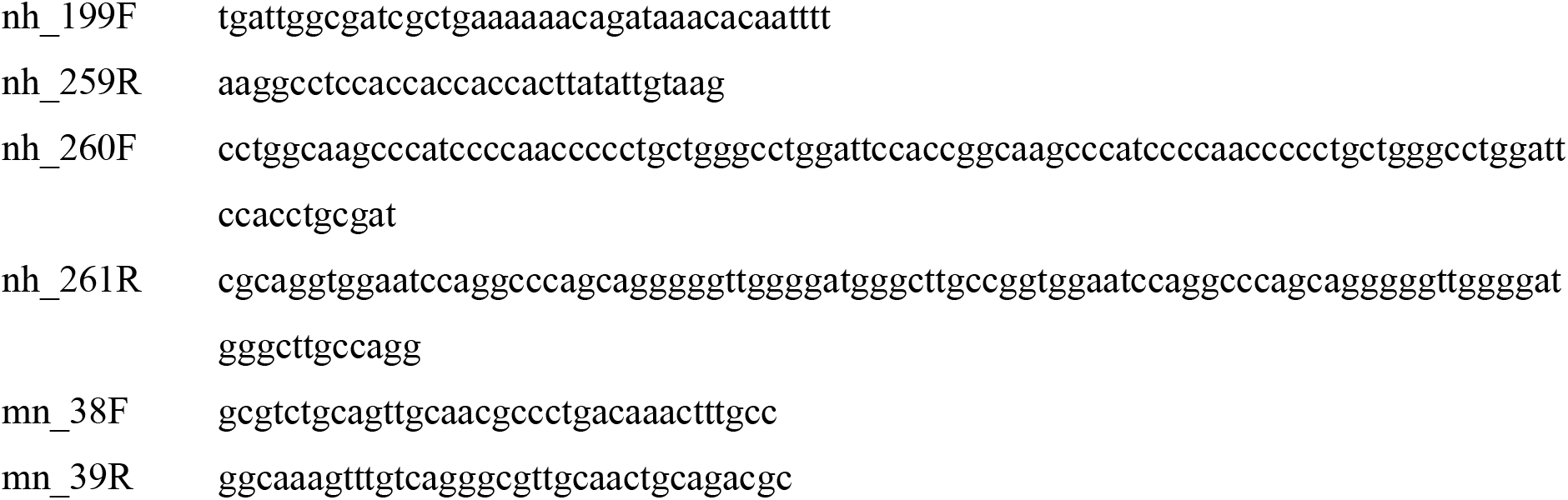

### PhiC31-mediated integration of transgenes

Plasmids containing *w+* and an *attB* site were injected into *phiC31[3xP3-RFP-3xP3-GFP-vas-PhiC31]; Cirl^KO^ attP-loxP;;* embryos (done by BestGene Inc.)^22^. *w+* served as the selection marker to identify recombinants and was subsequently removed by Cre-mediated excision (done by BestGene Inc.). Precise transgene insertion and removal of *w+ cassette* was validated by PCR genotyping.

### RT-PCR

Total RNA was isolated from larvae and adult *w^1118^* animals using RNeasy RNA Isolation Kit (Qiagen) and was directly used for reverse transcription using Superscript III Reverse Transcriptase according to manufacturer’s protocol (Invitrogen). PCR-amplification on transcribed cDNA libraries was carried out with the following primers:

#### Primers (5’-3’ orientation)

tl_5F: tcatcagggagcgcagcgtggtgca
tl_6R: atgctggtatagatcgaggtgcgcg
tl_392R: cgtgggtctggtcgcgaatattgt
rk_16: gtgaattttccttgtcgcgtg
rk_17: ctccagtctcgctgaagaag

### RNA-sequencing analyses

RNA-seq datasets that document the transcriptome at 30 distinct developmental stages of the *y1; cn bw1 sp1 Drosophila melanogaster* strain^34^ from NCBI Sequence Read Archive (SRA)^78^ were chosen to analyze and quantify *Cirl* splice variants during ontogenesis. Reads were mapped to the reference *Drosophila melanogaster* genome (July 2014, BDGP6 (GCA_000001215.4)) with Ensembl 88 annotations using Tophat 2.0.14.^79^, which aligns reads using Bowtie2 (version 2.1.0). Reads, which did not map uniquely to a genome position were excluded. After indexing with SAMtools (version 1.3.1)^80^, the mapped reads were assembled to transcripts and quantified by StringTie (version 1.3.3)^81,82^. For Tophat, we used the ‘default’ parameters, which are commonly used in most studies. StringTie parameters ‘read coverage’ (-c), ‘transcript length’ (-m) and ‘bases on both sides of a junction a spliced read has to cover’ (-a) were set to minimal values in order to avoid missing transcripts and generating a bias. The parameter ‘fraction of most abundant transcript at one locus’ (-f) was lowered from default (0.01) to 0 since correction for artifacts and incompletely processed mRNA with a 1% cutoff was performed after the comparative analysis. For all other StringTie parameters default values were used. Assembled transcripts were inspected with the Integrated Genome Viewer (Broad Institute) (version 2.3.91)^83,84^, and samples showing a visible 3′ bias due to oligo-dT/poly-A primer selection were not included.

### Cell culture

HEK293T cells (RRID:CVCL_0063) were cultivated in Dulbecco’s Modified Eagle Medium (DMEM; Gibco, #11995065) supplemented with 10% (v/v) fetal bovine serum (Sigma-Aldrich, #F7524 or Gibco, #10500) and 1% penicillin/streptomycin (Sigma-Aldrich, #P4333 or Gibco, #15140122) at 37 °C and a 5% CO_2_ humidified atmosphere.

HEK293T cells were split into 6-well cell culture plates (4×10^5^ cells/well) for co-immunoprecipitation analysis and into 96-well cell culture plates (3×10^4^ cells/well) coated with 0.01% poly-L-lysine (Sigma-Aldrich, #P9404) for ELISA. After 24 h, cells were transiently transfected with 2.5 μg plasmid DNA per well (6-well plate) for co-immunoprecipitation, 25 ng receptor plasmid and 75 ng empty vector plasmid per well (96-well plate) for ELISA, respectively, using Lipofectamine 2000 (Invitrogen, #11668019) according to manufacturer’s recommendations. Cells were incubated 48 h for co-immunoprecipitation and 24 h for the cell surface ELISA in a 5% CO_2_ humidified atmosphere at 37 °C. After 24 h, cell culture medium was exchanged against 2 ml DMEM for co-immunoprecipitation.

### ELISA

For total and cell surface expression, transfected cells (equimolar amounts of DNA) were fixed with 50 μl of 4% (w/v) paraformaldehyde in PBS at RT for 10 min. Next, unspecific epitopes were blocked with 100 μl phosphate buffered saline (PBS) containing 5% (v/v) goat serum (Sigma-Aldrich, #G6767) at RT for 1 h. Then, cells were incubated for 1 h at RT in a 1:1000 dilution of α-HA-peroxidase (RRID: AB_390917) in 100 μl PBS containing 5% (v/v) goat serum. After two washing steps using PBS, the cells were incubated with a substrate solution (1 mg/ml *o*-phenylenediamine and 1 μl/ml hydrogen peroxide solved in a 0.05 M citric acid and 0.05 M disodium phosphate solution; pH 5) for 5 – 10 min at RT. The reaction was stopped with 100 μl 2.5 M sulfuric acid. 150 μl of the supernatant was used for absorbance measurements at 490 nm with a multimode-reader (Molecular Devices, SpectraMax M). For total ELISA, PBT (PBS + 0.05% Triton-X100) was used instead of PBS throughout the experiment.

### Immunofluorescence and imaging

#### Drosophila larvae

Third instar *Drosophila* larvae were dissected in ice-cold HL-3 (hemolymph-like solution)^85^, fixed for 10 min using cold 4% paraformaldehyde in 0.1 M phosphate buffered saline (PBS) and blocked overnight in PBT (PBS with 1% Triton X-100, Sigma-Aldrich), containing 5% normal goat serum (NGS, Jackson ImmunoResearch). Primary antibody incubation was done at 4°C overnight. The next day, secondary antibody incubation was done for 24 h at 4 °C. Each antibody incubation step was followed by two short and three 20 min washing steps (PBS with 1 % Triton X-100). Samples were stored in Vectashield (Vector Laboratories) overnight at 4 °C before mounting. Confocal imaging was performed with a Leica SP8 confocal system. To ascertain comparability between different genotypes, larvae were imaged in one session and image acquisition of genotypes alternated. Image analysis was done using ImageJ (NIH). Antibodies and dyes were used at following dilutions: rabbit-α-dsRed (1:500, Takara, #632496; RRID:AB_10013483), α-HRP conjugated with Alexa Fluor-488 (1:250; Jackson Immuno Research, #123-545-021, RRID: 2338965), Cy5-conjugated goat-α-rabbit (1:250; Jackson Immuno Research, #111-175-144, RRID: AB_2338013).

### Protein Isolation

#### HEK293T cells

Cells were lysed using 200 μl M-PER buffer (ThermoFisher, #78503) supplemented with protease inhibitor (1:1000, Sigma-Aldrich, #P8340). After addition of lysis buffer, cells were detached from the culture dish bottom using a cell scraper and incubated on ice for 5 min. Transferred samples were then centrifuged for 5 min at 14,000 rpm (4 °C) and a sample of the supernatant diluted 4:1 with 4x sample buffer (Licor, #928-40004) containing 10% (v/v) β-mercaptoethanol. Total protein concentration was determined using the Pierce BCA Protein Assay Kit (ThermoFisher, #23227). Note that in intra-co-immunoprecipitation experiments the same amount of total protein was used for all samples.

#### Fly heads

Fly heads (25/genotype) were severed using fine scissors (Fine Science Tools, #15005-08), collected in a 0.5 ml Eppendorf tubes and immediately frozen in liquid nitrogen. Subsequently, the heads were homogenized in 2% SDS supplemented with a protease inhibitor cocktail (1:1000; Sigma-Aldrich) using a glass rod (4°C). Next, Triton X-100 was added (final concentration 1%) to the samples, which were then centrifuged for 15 min (14,000 rpm) at 4 °C. The supernatant was collected in fresh, pre-cooled tubes and centrifuged twice again for 30 min at 14,000 rpm and 4 °C. Finally, the supernatant was supplemented with SDS-based sample buffer (Li-cor) and β-mercaptoethanol (Merck).

#### Pupae

One- to two-day-old pupae (~ 0.6 g/genotype) were collected, immediately frozen in liquid nitrogen and homogenized in a pre-cooled mortar using a pestle. The resulting protein powder was transferred into a pre-cooled tube and supplemented with homogenization buffer (50 mM Tris, 150 mM NaCl, 1% Triton X-100, Protease Inhibitor 1:1000, 1 mM DTT, 0.5 mM PMSF). Samples were further homogenized, first using an Ultra-Turrax (4 x 15 sec; IKA T10 Basic) and second using a glass homogenisator (squished 20x/sample). Next, samples were centrifuged at 4,000 rpm for 30 min, again at 13,000 rpm for 30 min and finally at 25,000 rpm for 30 min. The samples were kept at 4 °C throughout the procedure. Finally, the supernatant was supplemented with Laemmli buffer.

### Pull down

Co-immunoprecipitation was performed using the Pierce Magnetic HA tag IP/Co-IP Kit (ThermoFisher, #88838). The manufacturer’s Manual IP/Co-IP protocol and the Elution Protocol using sample buffer were applied.

### Immunoblots

Protein samples were electrophoretically separated using 8%, 4-12% or 4-20% Tris-glycine gradient gels and blotted (protein from fly heads: 15 V, 7 min; protein from pupae: 20 V, 1 min; 23 V, 4 min; 25 V, 2 min; protein from HEK293T cells: 15 V, 6 min) onto nitrocellulose membrane (0.2 μm pore diameter, iBlotTM Transfer stack, ThermoFisher, #IB23001/#IB23002) using the iBlot2 (Invitrogen). The membrane was blocked for 1h using Odyssey Blocking buffer (Li-cor, #927-40000) diluted 1:2 with PBS. Blots were probed with primary antisera at indicated concentrations overnight at 4 °C: rabbit-α-RFP (1:1000, Antibodies-Online, RRID:AB_10781500), mouse-α-V5 (1:1000, Invitrogen, RRID:AB_2556564), rabbit-α-HA (1:1000, C29F4, Cell signaling technology; RRID:AB_1549585), rabbit-α-Tubulin*α* (1:5000, Santa Cruz antibodies, #12462-R), mouse-α-Tubulin*β* (1:2000, Developmental Studies Hybridoma Bank e7, RRID: AB_528499).

After rinsing twice and 3 x 5 min washing steps using 0.1% Tween-20-containg PBS, membranes were incubated with secondary antibodies from Li-cor (1:15000): IRDye 680RD goat-α-mouse (RRID: AB_10956588), IRDye 680RD goat-α-rabbit (RRID: AB_10956166), 800CW goat-α-mouse (RRID: AB_621842), 800CW goat-α-rabbit (RRID: AB_621843) for 1 h at RT and again rinsed twice and washed 3×5 min. Western blots were imaged with an OdysseyFc 2800 (Li-cor).

### Larval locomotion

Briefly, third instar larvae were positioned in the center of a 9-cm petri dish filled with 1% agarose (in ddH_2_O) and their behavior was recorded for two minutes using a camera (Logitech C920 HD Pro). The recorded videos were tracked and analyzed with a custom-written software (for details, see^86^). We calculated the speed of the midpoint of the larvae in mm/s. For the data displayed in Figure 3B, we calculated the average midpoint speed of all larvae on a given petri dish over 2 min – thus, the sample size represents the number of experimental repetitions. For the display in Figure 3A we binned the data in 5 s intervals, and calculated the average midpoint speed, as well as the 95% confidence interval of all data of a given genotype in each bin. Then, we smoothened the curve with a sliding average filter of 5 bins. Data were analyzed with Prism 7 (GraphPad).

### Ca^2+^ Imaging

To compare the effects of different *Cirl* mutations on lch5 function, the genetically encoded Ca^2+^ indicator jGCaMP7f^36^ was expressed under *UAS* control, using the *iav-gal4* driver^22^ in homozygous *Cirl^Rescue^*, *Cirl^KO^*, *Cirl^1TM^*, *Cirl^7TM^* and *Cirl^1TM+7TM^* backgrounds. Non-wandering male third instar larvae were pinned to a Sylgard pad in ice-cold, Ca^2+^ free HL-solution^22,87^ and cut open along the dorsal midline. The body muscle wall was gently stretched to the sides and pinned to the pad with fine dissection needles (#26002-10, Fine Science Tools). Innards and the CNS were gently removed. The Sylgard pad was transferred to the imaging chamber, filled with 2 mM CaCl2-containing HL-solution. To stimulate the lch5 mechanically, a 900-Hz tone (sine wave) of different intensities (75 – 90 dB SPL, starting with 75) was applied via a small suspended computer loud speaker. Lch5 activity pattern was unchanged even when stimulus direction was moved 180° from its initial position (data not shown). The sound frequency was controlled by Audacity software (V3.0.0, audacityteam.org). Stimulation intensity was verified by measuring sound pressure levels prior to every experiment. At small stimulation intensities, several measurements were averaged to increase the signal to noise ratio: 75 dB, 10 measurements; 80 dB, 5 measurements. Fluorescence imaging of the jGCaMP7f signal was performed on a DM 6FS (Leica) upright microscope equipped with a 40x, 0.8 N.A. dip-in objective and a GFP filter cube: excitation, 450 – 490 nm; dichroic 495 nm; emission, 500 – 550 nm. After pre-bleaching samples for 90 s to reduce bleaching rates during the experiment, samples were illuminated by blue LED light of the same intensity (LedHUB equipped with a LedH.465.4600, Omicron Laserage). jGCaMP7f fluorescence signals were video-recorded with a Hamamatsu Orca flash 4.0, V3 sCMOS camera at a 25-Hz frame rate and a 163 x 163 nm pixel size. Ten larvae of each genotype were measured. Videos were digitized and stored using the HOKAWO 3.0 software (Hamamatsu) and imported to Fiji ImageJ (NIH). A round, five-pixel diameter region of interest (ROI) was placed at the distal end of lch5 dendrites. The time course of the mean fluorescence intensity (F) in the ROI was exported to GraphPad Prism 6. ΔF/F_0_ was determined by calculating F_0_, i.e. the mean intensity during the 0.36 s period before stimulation and normalizing ΔF (i.e. F - F_0_) to F_0_. To quantitatively compare Ca^2+^ signals between genotypes, the ΔF/F_0_ averages of the Ca^2+^ signal plateaus (time 1.68 – 2.68 s) were calculated. Videos, in which any movement was detected were discarded.

### Electrophysiology

To compare the effects of different *Cirl* alleles on lch5 function, non-wandering male third instar larvae were dissected as described in ‘Ca^2+^ imaging’. After the innards and the CNS were removed, longitudinal muscles of the third, right abdominal hemi segment were cut cautiously. The lch5 nerve bundle was cut near the ventral, anterior end of muscle 21. The muscles 21, 22 and 23 were carefully cut and removed. The Sylgard pad was transferred to the recording chamber, filled with 2 mM CaCl2-containing HL-solution^88^. Micropipettes with 6 – 10 μm diameters were fabricated using the DMZ Universal Puller (Zeitz-Instrumente). To perform action current (AC) recordings, the pipette, which harbored the Ag/AgCl electrode, was filled with the bath solution. The lch5 nerve bundle was sucked into the pipette, and its opening was moved close to the soma of the neurons. Electrical activity was measured in voltage-clamp mode at 0 V using the Axopatch 200B amplifier (Molecular Devices). The measured current was low-pass Bessel filtered at 1 kHz and digitized at 10 kHz with the Axon Digidata 1550B analog-digital converter and Clampex 10.2 Software (both from Molecular Devices). To stimulate the lch5 neurons, cap cells were perpendicularly hooked with a miniature stimulation hook made from fine dissection needles (#26002-10, Fine Science Tools). The hook was attached to a piezo element (Physik Instrumente, P-840.30). Pulls of 0.03, 0.1, 0.3, 1, and 3 μm were administered by applying negative voltage jumps to the piezo using the E663 amplifier (Physik Instrumente), thus deflecting the lch5 to the side. Pull lengths of 3 μm and 1 μm were confirmed optically. The voltages for the 0.3, 0.1, and 0.03 μm had to be extrapolated from the voltages of longer pulls. Although the setup was thoroughly mechanically uncoupled, the hook most probably moved relatively to the organ by a much larger distance than the shortest pull length. Therefore, a reliable 30 nm pull seems unrealistic at first glance. However, since the piezo movement is very fast, its stability was measured as ±10 nm, and the organ reacts to stimulation in a differential manner, a meaningful reaction to the 30 nm pull could be easily detected. Each pull length, starting with 0.03 μm, was applied three times for 500 ms with a one-second break between the pulls. After a 7 s pause, the next pull length was applied three times. Action currents (AC), which occurred during the first 50 ms of the pull were manually counted. The AC frequency was calculated and evaluated in SigmaPlot 12 (SyStat). The mean of the three values of the same pull length was calculated to give n = 1. Ten measurements in ten different animals per genotype were performed. Treatment with the adenylyl cyclase inhibitor was carried out by adding 100 μM SQ22536 (Merck, #568500) to the bath solution and incubate it for 10 min before starting the electrophysiological measurement. For sham treatment, the adenylyl cyclase inhibitor was replaced with water.

### Statistics

If not stated otherwise, Data are reported as mean ± SEM, n indicates the sample size, p-values are given as an estimate of statistical significance. Data were analyzed with Prism 6 and 7 (GraphPad) and Sigma Plot 12 (Systat). If not stated otherwise, group means were compared by two-tailed Student’s t-tests, unless data was non-normally distributed. In this case, group means were compared by non-parametric tests, Mann-Whitney test for unpaired comparisons and Wilcoxon Signed Rank test for paired comparisons.

## ACKNOWLEDGEMENTS

We thank Maria Oppmann, Uta Strobel for technical assistance, Vladimir Katanaev for materials and Bertram Gerber and Diana Le Duc (D.L.D) for discussions. This work was supported by grants from the Deutsche Forschungsgemeinschaft to N.S. (265903901, FOR 2149/P01 [SCHO1791/1-2] and CRC 1423 project number 421152132, subproject B06); T.L. (265903901, FOR 2149/P01 and P03, SFB 1047/A05, TRR 166/C03, LA2861/7-1, CRC 1423 project number 421152132, subproject B06); T.S. (CRC 1423 project number 421152132, C04); MBK was partially funded through the Else-Kröner-Fresenius Stiftung (2020_EKEA.42 to D.L.D.) and N.S. and D.L. through Junior research grants from the Medical Faculty, Leipzig University.

## AUTHOR CONTRIBUTIONS

N.S. and D.L. conceived and supervised the study; prepared figures and wrote the manuscript with consent from all co-authors.

Conceptualization NS, DL, TL

Methodology NS, DL, MT, MS, MB, MBK, JI, TL, JL, MSG, JA

Validation NS, DL, MB, DCNH, MBK, AB, AKD

Formal analysis NS, DL, MT, MS, MB, AB, MBK, JI, AKD, TL, MSG

Investigation NS, DL, TS, VL, MB, AB, MBK, EK, JI, DCNH, AKD, TL, JL, MSG, JA

Resources

Writing – Original draft NS, DL

Writing – Review & Editing NS, DL, TL, TS, JL, AB, MBK, AKD, JL Visualization NS, DL, AB

Supervision NS, DL

Project administration NS, TL

Funding Acquisition NS, DL, TL, TS

## SUPPLEMENTAL INFORMATION

**Figure S1.**
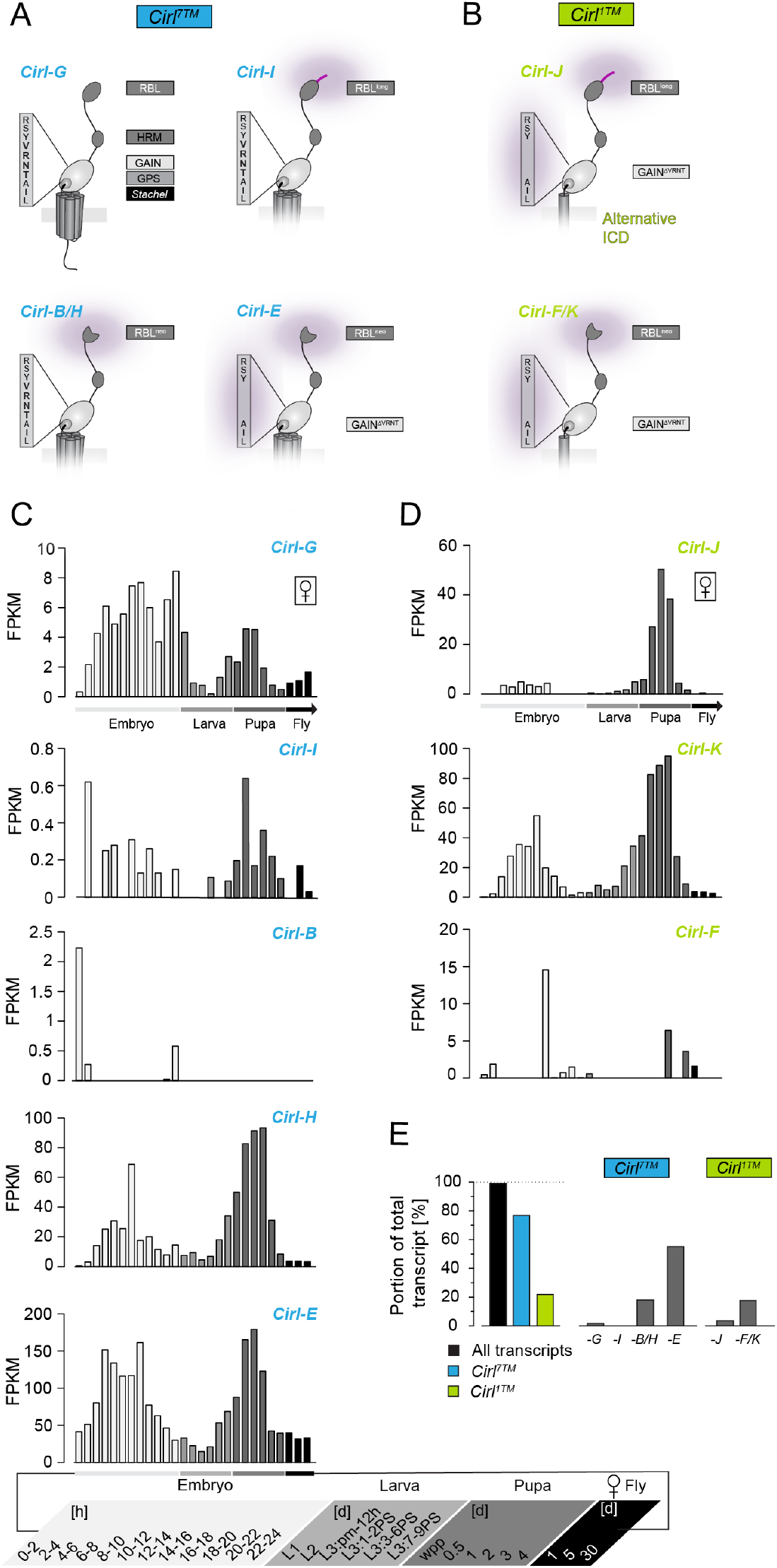
Ontogenetic analysis of the *Cirl* transcriptome. **(A, B)** Schematic of receptor isoforms encoded by the *Cirl* locus. Eight different *Cirl* transcripts are produced from a single locus. Five of them encode canonical receptors that contain 7TMs (indicated in blue) **(A)**, while the rest encodes non-canonical protein with only 1TM (indicated in green) **(B)**. *Cirl^7TM^* transcripts *-B* and *-H* as well as *Cirl^1TM^* transcripts *–K* and *–F* are translated into identical proteins, respectively, yielding six putative isoforms. Both species contain transcripts with a truncated N terminus (RBL^neo^ domain) and/or a four amino acid deletion in the GAIN domain (GAIN^ΔVRNT^). All transcripts contain an intact *Stachel* sequence ^41,89^. **(C, D)** Bioinformatic analyses of mRNAs isolated from *y1; cn bw1 sp1 Drosophila* strain at different developmental stages ^34^ revealed peak expression of *Cirl^7TM^* (left panel) and *Cirl^1TM^* (right panel) transcript in embryo and pupae. Embryos were collected every two hours for one day; L1 larvae were harvested 42-44 h after egg-laying, L2 larvae 66-70 h after egg-laying and L3 larvae 83-85 h after egg-laying; PS: puff stages; wpp: white prepupae^34^. **(E)** *Cirl^7TM^* and *Cirl^1TM^* transcript quantification shows that ~ 80% of *Cirl* mRNA encodes Cirl^7TM^ protein, ~ 74.7% of which present *Cirl-B/H* and *-E* transcripts. The remaining *Cirl* mRNAs encode Cirl^1TM^ proteins (*-F/K*: 18.3%; *-I*: 4.2%).

**Figure S2.**
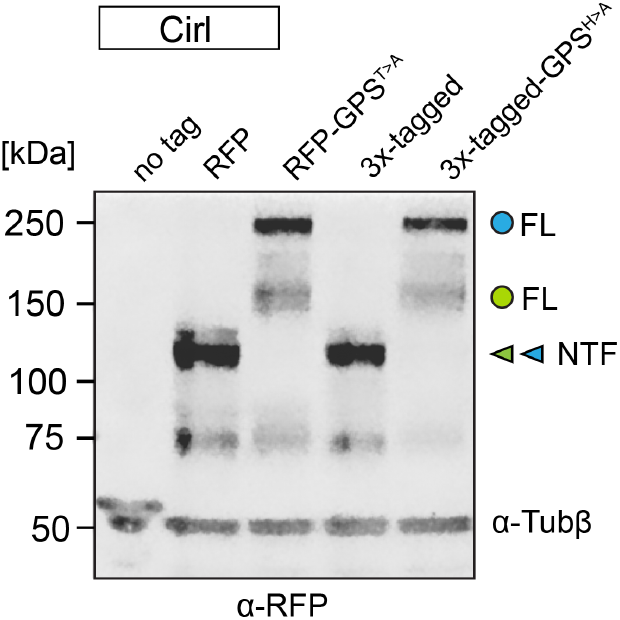
Cirl^1TM^ is expressed in the adult fly brain. Western blot of head homogenates shows expression of Cirl^7TM^ and Cirl^1TM^. An α-RFP antiserum recognized the NTFs of RFP∷Cirl^21^ and Cirl3x-tagged (mRFP also inserted in the NTF^21^; arrowheads green: Cirl^1TM^; blue: Cirl^7TM^) as well as the FL-1TM (indicated by green circle) and FL-7TM molecules of RFP∷Cirl^GPS/T>A^ and Cirl3x-tagged^GPS/H>A^ animals (indicated by the blue circle). An additional Cirl band of unknown identity was detected (~ 70 kDa). Tubulin served as a loading control (α-Tub*β*).

**Figure S3.**
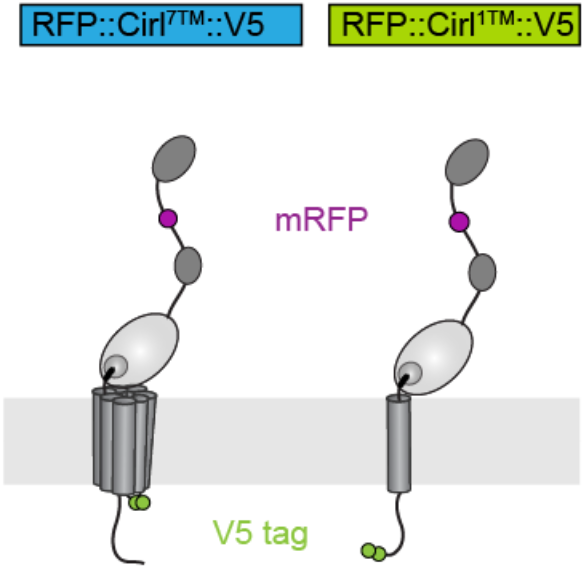
Scheme of RFP-tagged Cirl isoform. Both Cirl^7TM^ and Cirl^1TM^ proteins contain an N-terminal mRFP. Cirl^7TM^ contains a tandem V5-tag in the last intracellular loop (RFP∷Cirl^7TM^∷V5). Cirl^1TM^ contains a tandem V5-tag at the C terminus (RFP∷Cirl^1TM^∷V5). Related to Figure 2.

**Figure S4.**
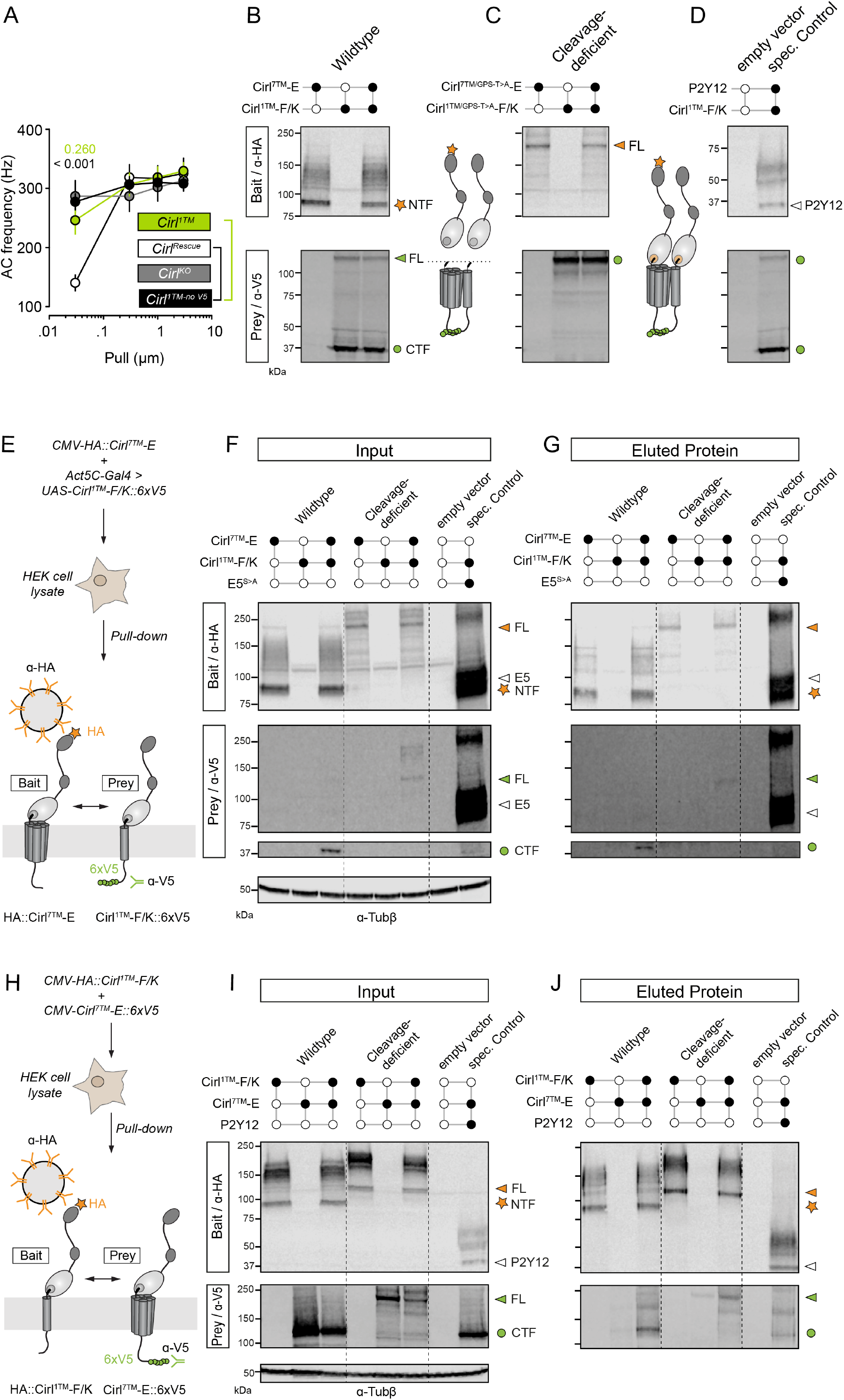
Cirl^1TM^ and Cirl^7TM^ co-immunoprecipitate in HEK cells when expressed using the UAS-GAL4 system. **(A)** Quantification of mechanically-evoked lch5 AC frequencies (pull lengths 0.03 - 3 μm). C-terminal addition of V5-tags has no impact on Cirl^1TM^ function. P-values denote statistical difference between *Cirl^1TM-no V5^* and *Cirl^Rescue^* (black) or *Cirl^1TM^* (green) at 30 nm pull length. Data are shown as mean ± SEM (each genotype n = 10). **(B-D)** Immunoblot input samples from the pull-down experiment shown in Figure 5E-G. Wildtype (B), cleavage-deficient Cirl (C) and Controls (D) were run on a gel side-by-side. Cirl^7TM^-E bait (orange star/arrowhead, upper row). Cirl^1TM^-F/K prey (green circle/arrowhead, lower row). HA-tagged P2Y12 served mock bait. Tubulin served as loading control (α-Tub*β*). **(E)** Schematic of the co-immunoprecipitation experiment shown in (F and G). Note that Cirl^1TM^-F/K∷6xV5 prey is expressed using *UAS-Gal4* binary expression system^42^. **(F, G)** Immunoblots of Cirl isoform input samples (F) and co-precipitation eluates (G). Cirl^7TM^-E served as bait (upper row); FL protein is indicated by orange arrow, NTF with an orange star. Cirl^1TM^-F/K prey (green circle/arrowhead, lower row). Cleavage-disabled HA-V5-tagged ADGRE5 served as mock bait (white arrowhead). Tubulin served as loading control (α-Tub*β*). **(H-J)** Immunoblots of Cirl isoform input samples (I) and co-precipitation eluates (J). Cirl^1TM^-F/K served as bait (upper row); FL protein is indicated by orange arrow, NTF with an orange star. Cirl^7TM^-E prey (CTF = green circle; FL = arrowhead, lower row). HA-tagged P2Y12 served as mock bait (white arrowhead). Tubulin served as loading control (α-Tub*β*).

**Figure S5.**
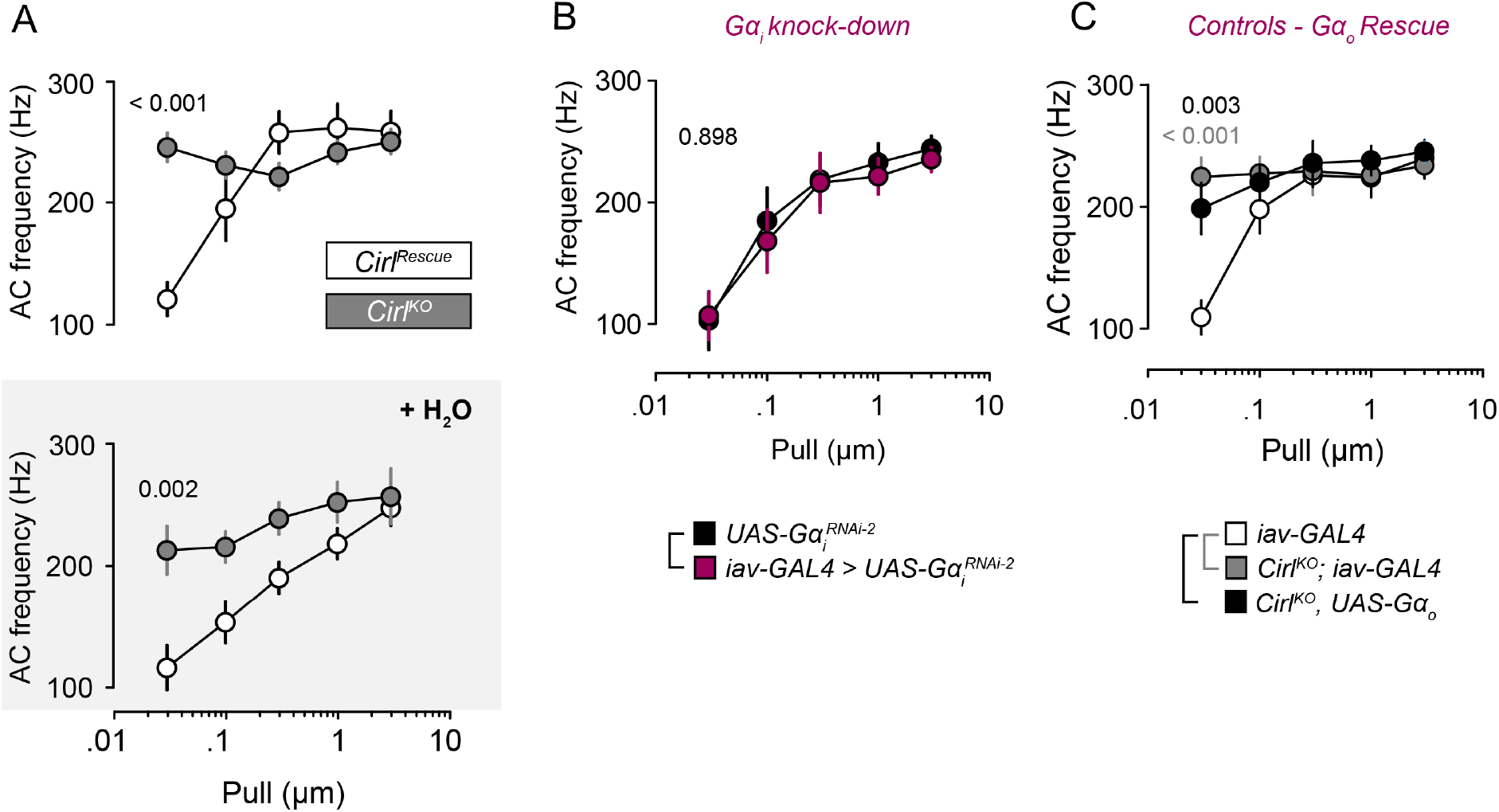
Sham treatment of *Cirl^KO^* lch5 leaves AC frequencies largely unaffected and overexpression of Gα_o_ protein rescues mechano-discrimination deficit of *Cirl^KO^*. **(A)** AC frequencies from mechanically stimulated lch5 neurons of *Cirl^KO^* animals before (A) and after sham treatment (+ H_2_O) show no effect. Related to Figure 6B-D and to Table S11. **(B)** AC frequencies from mechanically stimulated Gαi-deficient lch5 neurons (*iav-GAL4 > UAS-Gα_i_^RNAi-2^*). Lch5 neuron-specific *Gα_i_* knock-down leaves AC frequencies unaffected. P-value is derived from the statistical comparison of the AC frequencies (30 nm pull) between *iav-GAL4* > *UAS-Gα_i_^RNAi-2^* and *UAS-Gα_i_^RNAi-2^*. Related to Figure 6E and Table S14. **(C)** AC frequencies from mechanically stimulated lch5 neurons of *Cirl^KO^* with a copy of *iav-GAL4* (gray) or *UAS-G*α*o* (black). Neither the presence of UAS nor GAL4 transgenes alone reduces AC frequencies. P-values are derived from the statistical comparisons of the AC frequencies (30 nm pull) between *iav-GAL4* and *Cirl^KO^, UAS-G*α*o* (black) or *iav-GAL4* and *Cirl^KO^, iav-Gal4* (gray). Data are shown as mean ± SEM (each genotype n = 10). Related to Figure 6G and Table S15.

**Table S1.**
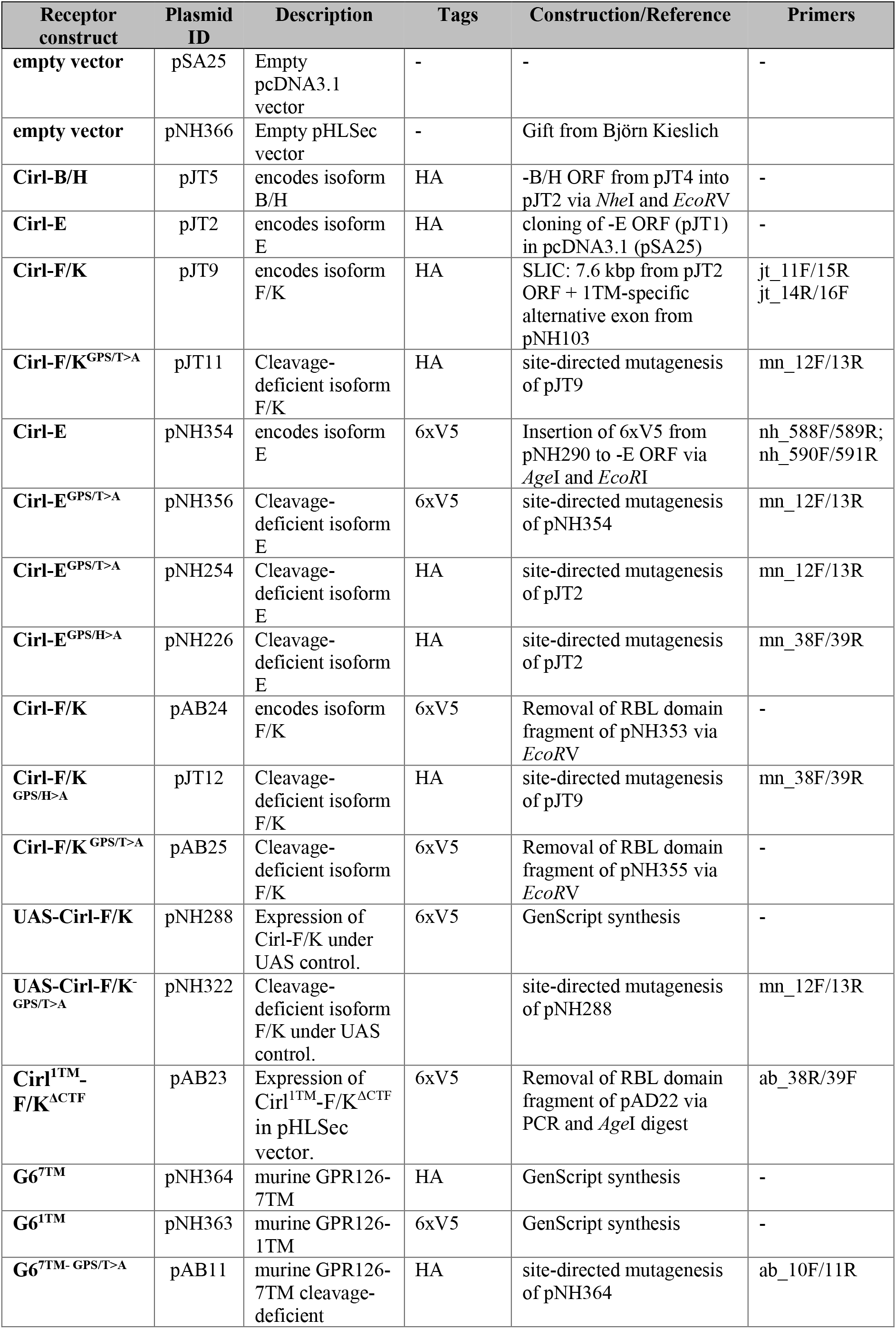
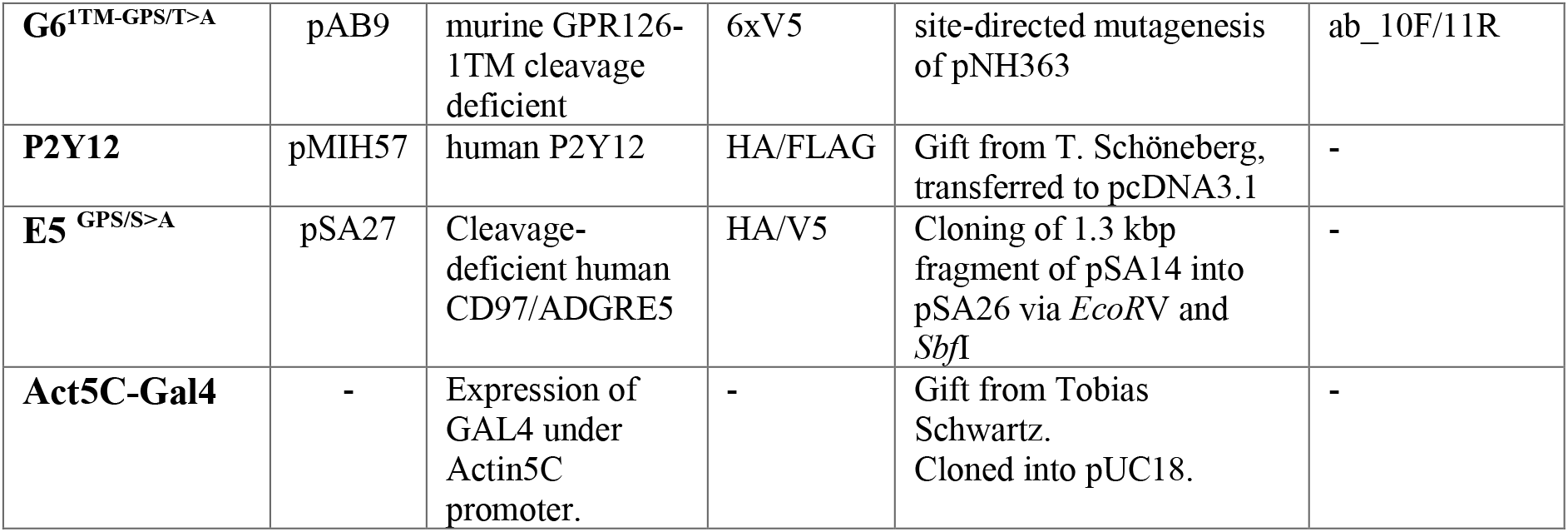
Collection of Cirl cDNA plasmids and their construction.

| Receptor construct | Plasmid ID | Description | Tags | Construction/Reference | Primers |
| --- | --- | --- | --- | --- | --- |
| empty vector | pSA25 | Empty pcDNA3.1 vector | - | - | - |
| empty vector | pNH366 | Empty pHLSec vector | - | Gift from Björn Kieslich |  |
| Cirl-B/H | pJT5 | encodes isoform B/H | HA | -B/H ORF from pJT4 into pJT2 via *NheI* and *EcoRV* | - |
| Cirl-E | pJT2 | encodes isoform E | HA | cloning of -E ORF (pJT1) in pcDNA3.1 (pSA25) | - |
| Cirl-F/K | pJT9 | encodes isoform F/K | HA | SLIC: 7.6 kbp from pJT2 ORF + 1TM-specific alternative exon from pNH103 | jt_11F/15R<br>jt_14R/16F |
| Cirl-F/K<sup>GPS/T>A</sup> | pJT11 | Cleavage-deficient isoform F/K | HA | site-directed mutagenesis of pJT9 | mn_12F/13R |
| Cirl-E | pNH354 | encodes isoform E | 6xV5 | Insertion of 6xV5 from pNH290 to -E ORF via *AgeI* and *EcoRI* | nh_588F/589R;<br>nh_590F/591R |
| Cirl-E<sup>GPS/T>A</sup> | pNH356 | Cleavage-deficient isoform E | 6xV5 | site-directed mutagenesis of pNH354 | mn_12F/13R |
| Cirl-E<sup>GPS/T>A</sup> | pNH254 | Cleavage-deficient isoform E | HA | site-directed mutagenesis of pJT2 | mn_12F/13R |
| Cirl-E<sup>GPS/H>A</sup> | pNH226 | Cleavage-deficient isoform E | HA | site-directed mutagenesis of pJT2 | mn_38F/39R |
| Cirl-F/K | pAB24 | encodes isoform F/K | 6xV5 | Removal of RBL domain fragment of pNH353 via *EcoRV* | - |
| Cirl-F/K<sup>GPS/H>A</sup> | pJT12 | Cleavage-deficient isoform F/K | HA | site-directed mutagenesis of pJT9 | mn_38F/39R |
| Cirl-F/K<sup>GPS/T>A</sup> | pAB25 | Cleavage-deficient isoform F/K | 6xV5 | Removal of RBL domain fragment of pNH355 via *EcoRV* | - |
| UAS-Cirl-F/K | pNH288 | Expression of Cirl-F/K under UAS control. | 6xV5 | GenScript synthesis | - |
| UAS-Cirl-F/K<sup>GPS/T>A</sup> | pNH322 | Cleavage-deficient isoform F/K under UAS control. |  | site-directed mutagenesis of pNH288 | mn_12F/13R |
| Cirl<sup>1TM</sup>-F/K<sup>ΔCTF</sup> | pAB23 | Expression of Cirl<sup>1TM</sup>-F/K<sup>ΔCTF</sup> in pHLSec vector. | 6xV5 | Removal of RBL domain fragment of pAD22 via PCR and *AgeI* digest | ab_38R/39F |
| G6<sup>7TM</sup> | pNH364 | murine GPR126-7TM | HA | GenScript synthesis | - |
| G6<sup>1TM</sup> | pNH363 | murine GPR126-1TM | 6xV5 | GenScript synthesis | - |
| G6<sup>7TM</sup>-GPS/T>A | pAB11 | murine GPR126-7TM cleavage-deficient | HA | site-directed mutagenesis of pNH364 | ab_10F/11R |
| **G6<sup>1TM-GPS/T>A</sup>** | pAB9 | murine GPR126-1TM cleavage deficient | 6xV5 | site-directed mutagenesis of pNH363 | ab_10F/11R |
| **P2Y12** | pMIH57 | human P2Y12 | HA/FLAG | Gift from T. Schöneberg, transferred to pcDNA3.1 | - |
| **E5<sup>GPS/S>A</sup>** | pSA27 | Cleavage-deficient human CD97/ADGRE5 | HA/V5 | Cloning of 1.3 kbp fragment of pSA14 into pSA26 via *EcoRV* and *SbfI* | - |
| **Act5C-Gal4** | - | Expression of GAL4 under Actin5C promoter. | - | Gift from Tobias Schwartz. Cloned into pUC18. | - |

**Table S2.**
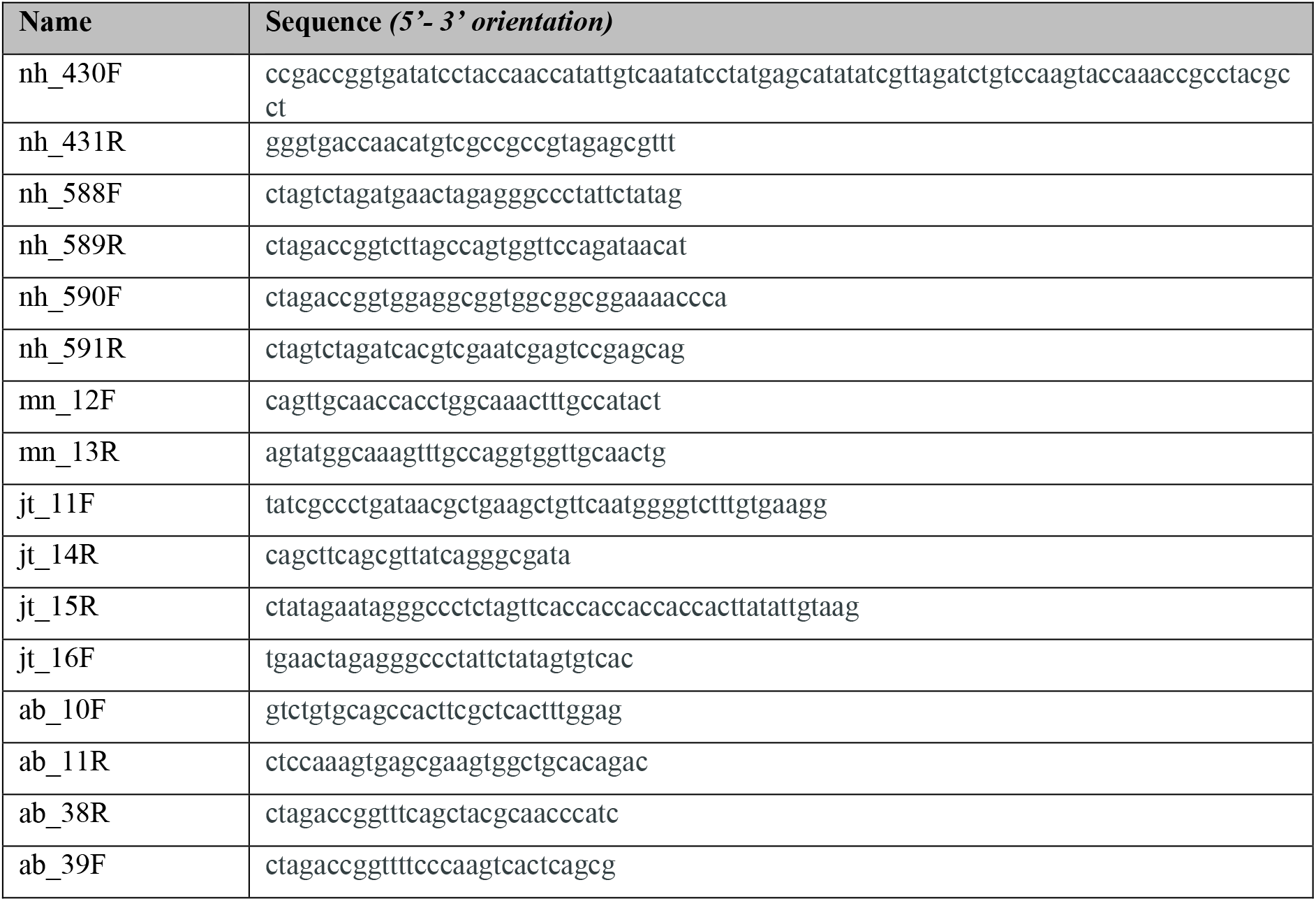
Collection of primers used for cloning of cDNA constructs.

**Table S3.** Quantification of mean midpoint speed (N = 20, n=5/dish). Values represent the mean ± SEM, N denotes number of experiments. All groups were compared against *Cirl^Rescue^* using an unpaired t-test.

| Genotype | Mean midpoint speed (mm/s) | N | p-value |
| --- | --- | --- | --- |
| Related to Figure 3B |  |  |  |
| *Cir1<sup>Rescue</sup>* | 0.90 ± 0.06 | 20 | - |
| *Cir1<sup>KO</sup>* | 0.61 ± 0.02 | 20 | < 0.001 |
| *Cir1<sup>7TM</sup>* | 0.94 ± 0.05 | 20 | 0.670 |
| *Cir1<sup>1TM</sup>* | 1.14 ± 0.05 | 20 | 0.004 |

**Table S4.**
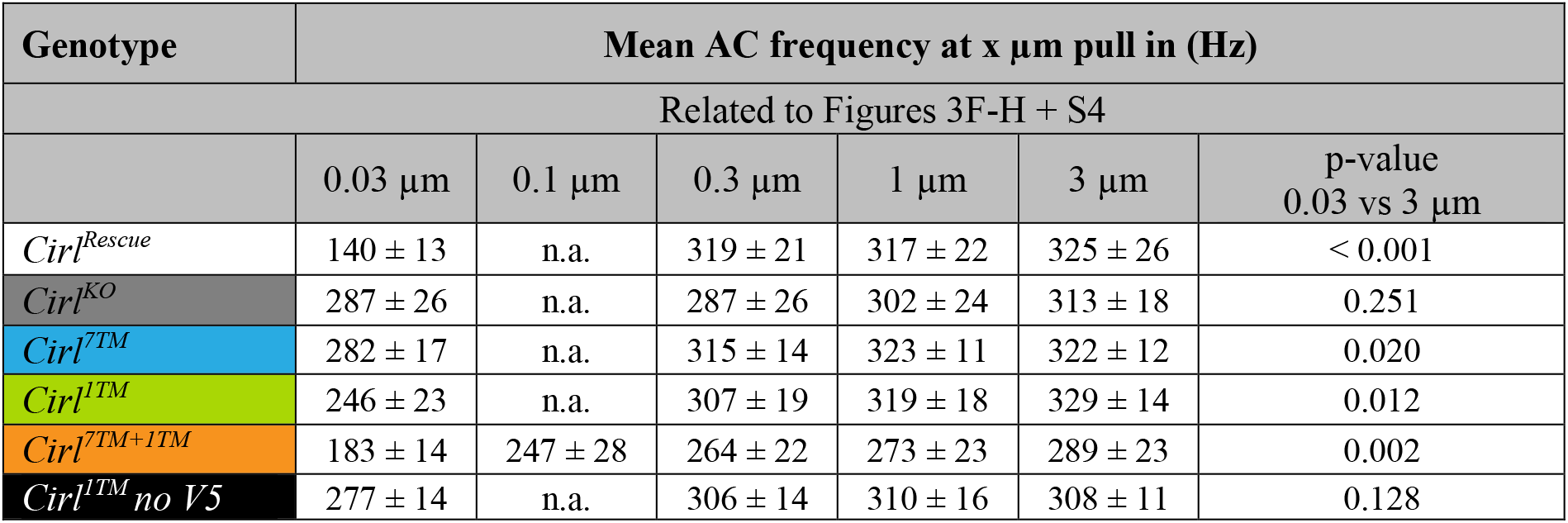
Quantification of AC frequencies of lch5 neurons at different pull lengths (0.03-3 μm). Values represent the mean ± SEM response of 10 different lch5 organs from 10 individual animals. AC frequencies at different pull lengths were compared using paired t-tests.

**Table S5.**
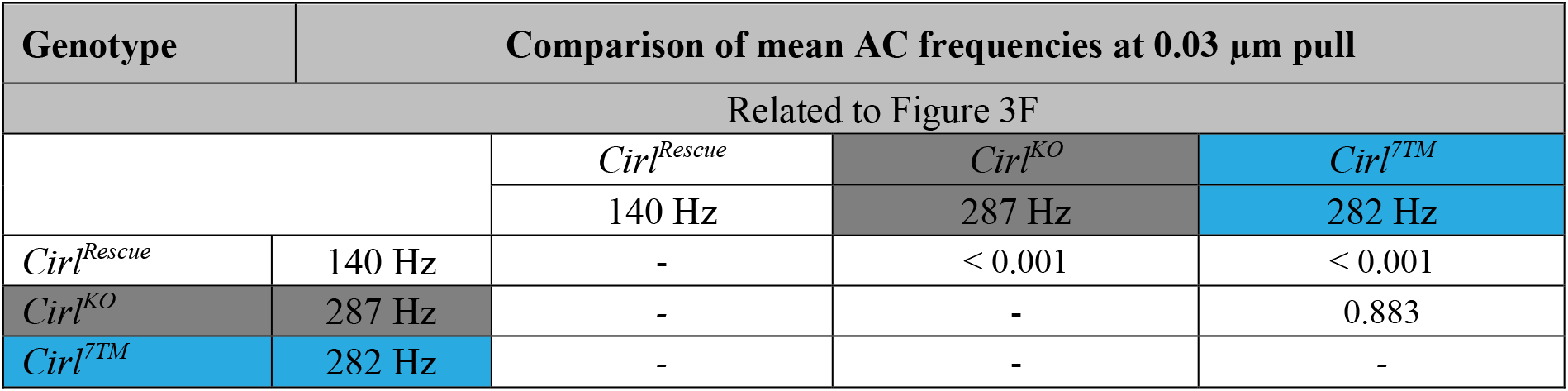
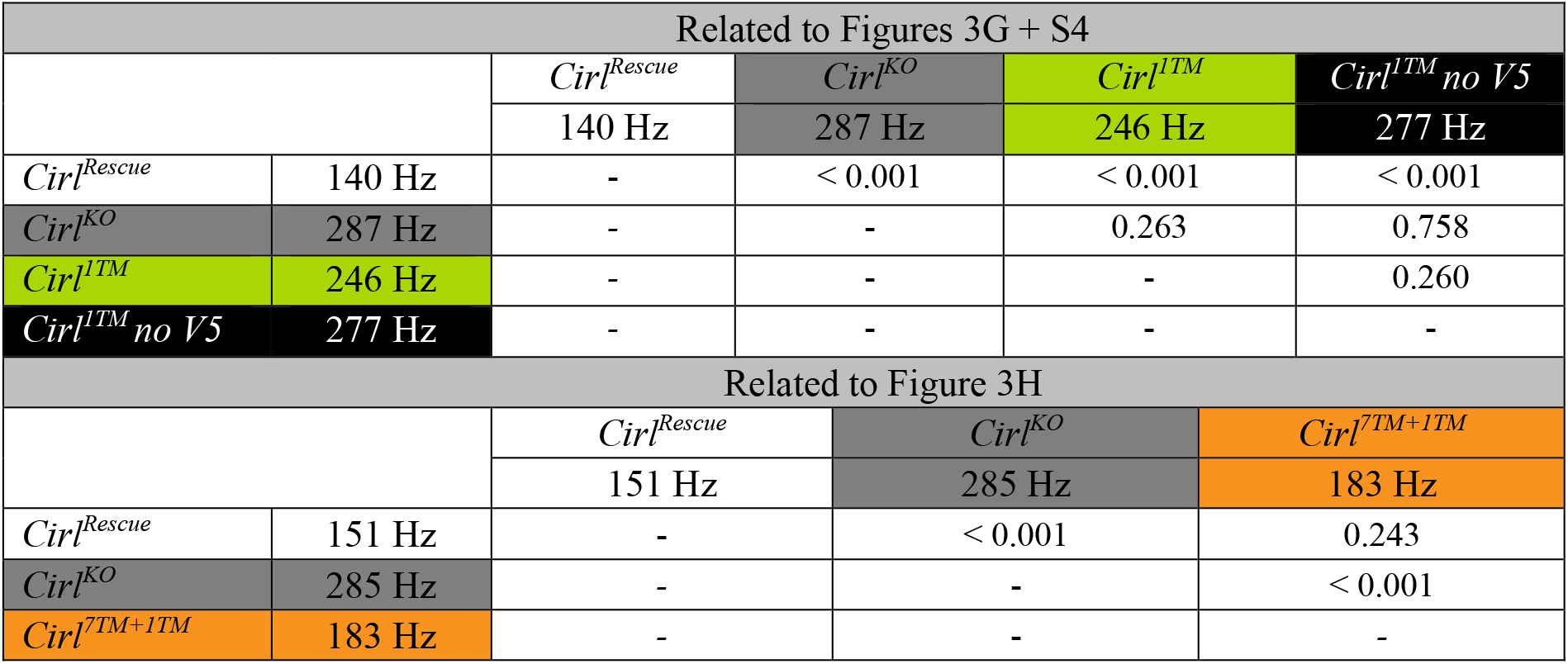
Comparison of AC frequencies generated in response to 0.03 μm pull. Values represent the mean response of 10 different lch5 organs from 10 individual animals. All groups were compared using unpaired t-tests.

| Genotype | Comparison of mean AC frequencies at 0.03 μm pull |  |  |  |
| --- | --- | --- | --- | --- |
| Related to Figure 3F |  |  |  |  |
|  |  | *Cir1<sup>Rescue</sup>* | *Cir1<sup>KO</sup>* | *Cir1<sup>7TM</sup>* |
|  |  | 140 Hz | 287 Hz | 282 Hz |
| *Cir1<sup>Rescue</sup>* | 140 Hz | - | < 0.001 | < 0.001 |
| *Cir1<sup>KO</sup>* | 287 Hz | - | - | 0.883 |
| *Cir1<sup>7TM</sup>* | 282 Hz | - | - | - |

| Related to Figures 3G + S4 |  |  |  |  |  |
| --- | --- | --- | --- | --- | --- |
|  |  | *Cirl<sup>Rescue</sup>* | *Cirl<sup>KO</sup>* | *Cirl<sup>1TM</sup>* | *Cirl<sup>1TM</sup> no V5* |
|  |  | 140 Hz | 287 Hz | 246 Hz | 277 Hz |
| *Cirl<sup>Rescue</sup>* | 140 Hz | - | < 0.001 | < 0.001 | < 0.001 |
| *Cirl<sup>KO</sup>* | 287 Hz | - | - | 0.263 | 0.758 |
| *Cirl<sup>1TM</sup>* | 246 Hz | - | - | - | 0.260 |
| *Cirl<sup>1TM</sup> no V5* | 277 Hz | - | - | - | - |
| Related to Figure 3H |  |  |  |  |  |
|  |  | *Cirl<sup>Rescue</sup>* | *Cirl<sup>KO</sup>* | *Cirl<sup>7TM+1TM</sup>* |  |
|  |  | 151 Hz | 285 Hz | 183 Hz |  |
| *Cirl<sup>Rescue</sup>* | 151 Hz | - | < 0.001 | 0.243 |  |
| *Cirl<sup>KO</sup>* | 285 Hz | - | - | < 0.001 |  |
| *Cirl<sup>7TM+1TM</sup>* | 183 Hz | - | - | - |  |

**Table S6.** Quantification of Ca^2+^ signals from the five larval lch5 neurons of *Cirl^Rescue^* at different sound-pressure levels (75 – 90 dB). Values represent the mean ± SEM response of different neurons from N individual animals. Ca^2+^ signals below ΔF/F_0_ < 0.01 are not shown.

| Ca<sup>2+</sup> signal as $\Delta F/F_0$ | | | | | | | | | | |
| --- | --- | --- | --- | --- | --- | --- | --- | --- | --- | --- |
| Related to Figure 4C |  |  |  |  |  |  |  |  |  |  |
| *Cir1<sup>Rescue</sup>* | Neuron 1 |  | Neuron 2 |  | Neuron 3 |  | Neuron 4 |  | Neuron 5 |  |
| | $\Delta F/F_0$ | N | $\Delta F/F_0$ | N | $\Delta F/F_0$ | N | $\Delta F/F_0$ | N | $\Delta F/F_0$ | N |
| 75 dB | 0.03 ± 0.01 | 10 | 0.05 ± 0.02 | 4 | < 0.01 | - | 0.05 ± 0.05 | 5 | < 0.01 | - |
| 80 dB | 0.09 ± 0.03 | 10 | 0.10 ± 0.03 | 6 | 0.01 ± 0.01 | 8 | 0.07 ± 0.04 | 5 | < 0.01 | - |
| 85 dB | 0.18 ± 0.03 | 10 | 0.22 ± 0.06 | 6 | 0.04 ± 0.02 | 8 | 0.14 ± 0.05 | 5 | < 0.01 | - |
| 90 dB | 0.73 ± 0.09 | 10 | 0.51 ± 0.10 | 6 | 0.07 ± 0.02 | 8 | 0.43 ± 0.13 | 5 | < 0.01 | - |

**Table S7.** Quantification of Ca^2+^ signals from neuron 1 of larval lch5 in different genotypes, at different sound-pressure levels (75 – 90 dB). Values represent mean responses ± SEM of 10 lch5 organs from 10 individual animals. Mean ΔF/F_0_ of *Cirl^Rescue^* and mutant genotypes at 75 dB were compared using unpaired t-tests, unless stated otherwise (*).

| Neuron 1 | Ca<sup>2+</sup> signal as ΔF/F<sub>0</sub> |  |  |  |  |
| --- | --- | --- | --- | --- | --- |
| Related to Figure 4D,E,F |  |  |  |  |  |
| Genotype | 75 dB | p-value | 80 dB | 85 dB | 90 dB |
| *Cir1<sup>Rescue</sup>* | 0.03 ± 0.01 | - | 0.09 ± 0.03 | 0.18 ± 0.03 | 0.73 ± 0.09 |
| *Cir1<sup>KO</sup>* | 0.17 ± 0.05 | 0.008 | 0.26 ± 0.06 | 0.40 ± 0.07 | 1.17 ± 0.20 |
| *Cir1<sup>7TM</sup>* | 0.15 ± 0.05 | 0.064* | 0.28 ± 0.07 | 0.48 ± 0.08 | 1.26 ± 0.19 |
| *Cir1<sup>1TM</sup>* | 0.13 ± 0.04 | 0.018 | 0.23 ± 0.04 | 0.42 ± 0.05 | 1.11 ± 0.20 |
| *Cir1<sup>7TM+1TM</sup>* | 0.10 ± 0.03 | 0.035 | 0.17 ± 0.05 | 0.28 ± 0.06 | 1.01 ± 0.17 |
| *Mann-Whitney test |  |  |  |  |  |

**Table S8.** Quantification of Ca^2+^ signals from lch5 neurons 2 and 4 in *Cirl^Rescue^* and *Cirl^KO^* larvae at different sound-pressure levels (75 – 90 dB). Values represent the mean ± SEM response of different neurons from N individual animals. Mean ΔF/F_0_ of *Cirl^Rescue^* and *Cirl^KO^* at 75 dB were compared using unpaired t-tests, unless stated otherwise (*).

|  | Ca<sup>2+</sup> signal as ΔF/F<sub>0</sub> |  |  |  |  |  |  |  |  |
| --- | --- | --- | --- | --- | --- | --- | --- | --- | --- |
| Related to Figure 4G,H |  |  |  |  |  |  |  |  |  |
| Neuron 2 | 75 dB | N | p-value | 80 dB | N | 85 dB | N | 90 dB | N |
| Cir1<sup>Rescue</sup> | 0.05 ± 0.02 | 4 | - | 0.10 ± 0.03 | 6 | 0.22 ± 0.07 | 6 | 0.51 ± 0.10 | 6 |
| Cir1<sup>KO</sup> | 0.14 ± 0.05 | 7 | 0.186 | 0.23 ± 0.05 | 6 | 0.19 ± 0.05 | 7 | 0.74 ± 0.16 | 7 |
| Neuron 4 |  |  |  |  |  |  |  |  |  |
| Cir1<sup>Rescue</sup> | 0.05 ± 0.05 | 5 | - | 0.07 ± 0.04 | 5 | 0.14 ± 0.05 | 5 | 0.43 ± 0.13 | 5 |
| Cir1<sup>KO</sup> | 0.09 ± 0.02 | 5 | 0.151* | 0.11 ± 0.03 | 5 | 0.11 ± 0.03 | 5 | 0.46 ± 0.09 | 5 |
| *Mann-Whitney test |  |  |  |  |  |  |  |  |  |

**Table S9.** Quantification of expression of certain Cirl isoforms via ELISA. Optical density was measured at 490 nm. P-values derived from statistical comparison to empty vector sample. Data represent the mean ± SEM (N=3, n=8) and were compared using unpaired t-tests, unless stated otherwise (*).

| Genotype | Cell surface expression | n | p-value |
| --- | --- | --- | --- |
| Related to Figure 5A |  |  |  |
| *Empty vector/pcDNA3.1* | 0.043 ± 0.001 | 24 | - |
| *HA::Cirl-B/H* | 0.36 ± 0.020 | 24 | < 0.001 |
| *HA::Cirl-E* | 0.32 ± 0.018 | 24 | < 0.001 |
| *HA::Cirl-F/K* | 0.63 ± 0.022 | 24 | < 0.001 |

| Genotype | Total expression | n | p-value |
| --- | --- | --- | --- |
| Related to Figure 5A |  |  |  |
| *Empty vector/pcDNA3.1* | 0.052 ± 0.001 | 24 | - |
| *HA::Cirl-B/H* | 0.59 ± 0.048 | 24 | < 0.001* |
| *HA::Cirl-E* | 0.32 ± 0.042 | 24 | < 0.001* |
| *HA::Cirl-F/K* | 0.70 ± 0.041 | 24 | < 0.001* |

\*Mann-Whitney test

| Genotype | Cell surface expression | n | p-value |
| --- | --- | --- | --- |
| Related to Figure 5C |  |  |  |
| *HA::Cirl-E* | 0.196 ± 0.011 | 24 | - |
| *HA::Cirl-E<sup>GPS-H>A</sup>* | 0.281 ± 0.010 | 24 | < 0.001 |
| *HA::Cirl-E<sup>GPS-T>A</sup>* | 0.426 ± 0.017 | 24 | < 0.001* |
| Related to Figure 5C |  |  |  |
| *HA::Cirl-F/K* | 0.570 ± 0.017 | 24 | - |
| *HA::Cirl-F/K<sup>GPS-H>A</sup>* | 0.525 ± 0.017 | 24 | 0.070 |
| *HA::Cirl-F/K<sup>GPS-T>A</sup>* | 0.593 ± 0.017 | 24 | 0.200* |

\*Mann-Whitney test

| Genotype | Total expression | n | p-value |
| --- | --- | --- | --- |
| Related to Figure 5C |  |  |  |
| *HA::Cirl-E* | 0.376 ± 0.020 | 24 | - |
| *HA::Cirl-E<sup>GPS-H>A</sup>* | 0.495 ± 0.019 | 24 | < 0.001 |
| *HA::Cirl-E<sup>GPS-T>A</sup>* | 0.606 ± 0.020 | 24 | < 0.001 |
| Related to Figure 5C |  |  |  |
| *HA::Cirl-F/K* | 0.664 ± 0.018 | 24 | - |
| *HA::Cirl-F/K<sup>GPS-H>A</sup>* | 0.726 ± 0.011 | 24 | < 0.001* |
| *HA::Cirl-F/K<sup>GPS-T>A</sup>* | 0.721 ± 0.017 | 24 | 0.008* |

\*Mann-Whitney test

**Table S10.** Pharmacological inhibition of ACyc. Quantification of lch5 AC frequencies at different pull lengths (0.03 - 3 μm) before and after a 10 min treatment with 100 μM adenylyl cyclase inhibitor SQ22536. Values represent mean responses ± SEM of 10 different lch5 organs from 10 individual animals. Mean frequencies of mutant genotypes at 0.03 μm pulls were compared to mean frequencies at the same pull length in *Cirl^Rescue^* using unpaired t-tests.

| Genotype | Mean AC frequency at x pull in (Hz) |  |  |  |  |  |
| --- | --- | --- | --- | --- | --- | --- |
| Related to Figure 6B-D |  |  |  |  |  |  |
|  | 0.03 μm | p-value | 0.1 μm | 0.3 μm | 1 μm | 3 μm |
|  | 0 min |  |  |  |  |  |
| *Cir1<sup>Rescue</sup>* | 133 ± 14 | - | 207 ± 27 | 269 ± 17 | 273 ± 19 | 270 ± 17 |
| *Cir1<sup>KO</sup>* | 257 ± 20 | < 0.001 | 245 ± 13 | 257 ± 15 | 269 ± 11 | 273 ± 14 |
| *Cir1<sup>7TM</sup>* | 270 ± 9 | < 0.001 | 245 ± 16 | 251 ± 11 | 259 ± 14 | 273 ± 15 |
| *Cir1<sup>1TM</sup>* | 270 ± 14 | < 0.001 | 263 ± 13 | 263 ± 13 | 259 ± 11 | 261 ± 9 |
|  | 10 min after addition of adenylyl cyclase inhibitor |  |  |  |  |  |
| *Cir1<sup>Rescue</sup>* | 139 ± 18 | - | 176 ± 17 | 213 ± 13 | 241 ± 13 | 270 ± 14 |
| *Cir1<sup>KO</sup>* | 139 ± 8 | 0.973 | 193 ± 11 | 230 ± 10 | 257 ± 9 | 286 ± 11 |
| *Cir1<sup>7TM</sup>* | 158 ± 10 | 0.366 | 175 ± 16 | 219 ± 22 | 242 ± 24 | 259 ± 16 |
| *Cir1<sup>1TM</sup>* | 175 ± 21 | 0.209 | 201 ± 17 | 215 ± 22 | 239 ± 20 | 271 ± 15 |

**Table S11.** Control experiment for pharmacological inhibition of ACyc. Quantification of lch5 AC frequencies at different pull lengths (0.03 - 3 μm) before and after a 10 min sham treatment to exclude an unspecific effect of the 10 min incubation. Values represent mean responses ± SEM of 10 lch5 organs from 10 individual animals. Mean frequencies of *Cirl^KO^* and *Cirl^Rescue^* at 0.03 μm pulls before and after treatment (sham treatment and 100 μM adenylyl cyclase inhibitor SQ22536 treatment, respectively) were compared using unpaired t-tests. Mean frequencies of *Cirl^KO^* at 0.03 μm pulls before and after sham treatment were due to non-normal data distribution compared using a Wilcoxon Signed Rank test (*).

| Genotype | Mean AC frequency at X pull in (Hz) |  |  |  |  |  |  |
| --- | --- | --- | --- | --- | --- | --- | --- |
| Related to Figure S5A |  |  |  |  |  |  |  |
|  | 0.03 μm | p-value<br>*Cirt*<sup>KO</sup> vs.<br>*Cirt*<sup>Rescue</sup> | 0.1 μm | 0.3 μm | 1 μm | 3 μm | p-value<br>0.03 μm<br>t<sub>0</sub> vs. t<sub>10min</sub> |
|  | 0 min |  |  |  |  |  |  |
| *Cirt*<sup>KO</sup> | 257 ± 12 | < 0.001 | 243 ± 11 | 233 ± 11 | 253 ± 9 | 262 ± 10 | - |
|  | 10 min after adding sham control |  |  |  |  |  |  |
| *Cirt*<sup>KO</sup> | 235 ± 20 | 0.002 | 238 ± 13 | 261 ± 13 | 275 ± 16 | 279 ± 23 | 0.652 |

**Table S12.** Mean AC frequency at different pull lengths in G protein knock-down. Quantification of AC frequencies of lch5 neurons at different pull lengths (0.03-3 μm). Values represent the mean ± SEM response of 10 lch5 organs from 10 individual animals. AC frequencies at different pull lengths were compared using paired t-tests.

| Genotype | Mean AC frequency at x $\mu\text{m}$ pull in (Hz) | | | | | |
| --- | --- | --- | --- | --- | --- | --- |
| Related to Figure 6E, F |  |  |  |  |  |  |
| | 0.03 $\mu\text{m}$ | 0.1 $\mu\text{m}$ | 0.3 $\mu\text{m}$ | 1 $\mu\text{m}$ | 3 $\mu\text{m}$ | p-value<br>0.03 vs 3 $\mu\text{m}$ |
| *iav-Gal4* | 121 ± 16 | 179 ± 32 | 235 ± 27 | 287 ± 22 | 279 ± 16 | < 0.001 |
| *UAS-G$\alpha_o$<sup>RNAi</sup>* | 114 ± 13 | 211 ± 27 | 228 ± 20 | 242 ± 12 | 248 ± 7 | < 0.001 |
| *UAS-Gα<sub>i</sub><sup>RNAi</sup>* | 132 ± 17 | 206 ± 24 | 231 ± 20 | 239 ± 15 | 249 ± 9 | < 0.001 |
| *iav-Gal4>UAS-Gα<sub>o</sub><sup>RNAi</sup>* | 242 ± 22 | 254 ± 23 | 245 ± 19 | 247 ± 14 | 249 ± 13 | 0.755 |
| *iav-Gal4>UAS-Gα<sub>i</sub><sup>RNAi</sup>* | 125 ± 20 | 227 ± 30 | 243 ± 30 | 256 ± 22 | 263 ± 12 | < 0.001 |

**Table S13.** Mean AC frequencies at 0.03 μm pull lengths in G protein knock-down. Comparison of AC frequencies generated by a 0.03 μm pull. Values represent the mean response of 10 lch5 organs from 10 individual animals. Unless stated otherwise (*), all groups were compared using unpaired t-tests.

| Genotype | | Comparison of mean AC frequencies at 0.03 μm pull | | | | |
| --- | --- | --- | --- | --- | --- | --- |
| Related to Figure 6E, F |  |  |  |  |  |  |
|  |  | *iav-Gal4/+* | *UAS-G$\alpha_o$<sup>RNAi</sup>* | *UAS-G$\alpha_i$<sup>RNAi</sup>* | *iav-Gal4>UAS-G$\alpha_o$<sup>RNAi</sup>* | *iav-Gal4>UAS-G$\alpha_i$<sup>RNAi</sup>* |
|  |  | 121 Hz | 114 Hz | 132 Hz | 242 Hz | 125 Hz |
| *iav-Gal4* | 121 Hz | - | 0.747 | 0.638 | < 0.001 | 0.858 |
| *UAS-G$\alpha_o$<sup>RNAi</sup>* | 114 Hz | - | - | 0.495* | < 0.001 | 0.638 |
| *UAS-G$\alpha_i$<sup>RNAi</sup>* | 132 Hz | - | - | - | < 0.001 | 0.805 |
| *iav-Gal4>UAS-G$\alpha_o$<sup>RNAi</sup>* | 242 Hz | - | - | - | - | < 0.001 |
| *iav-Gal4>UAS-G$\alpha_i$<sup>RNAi</sup>* | 125 Hz | - | - | - | - | - |
| *Mann-Whitney test |  |  |  |  |  |  |

**Table S14.**
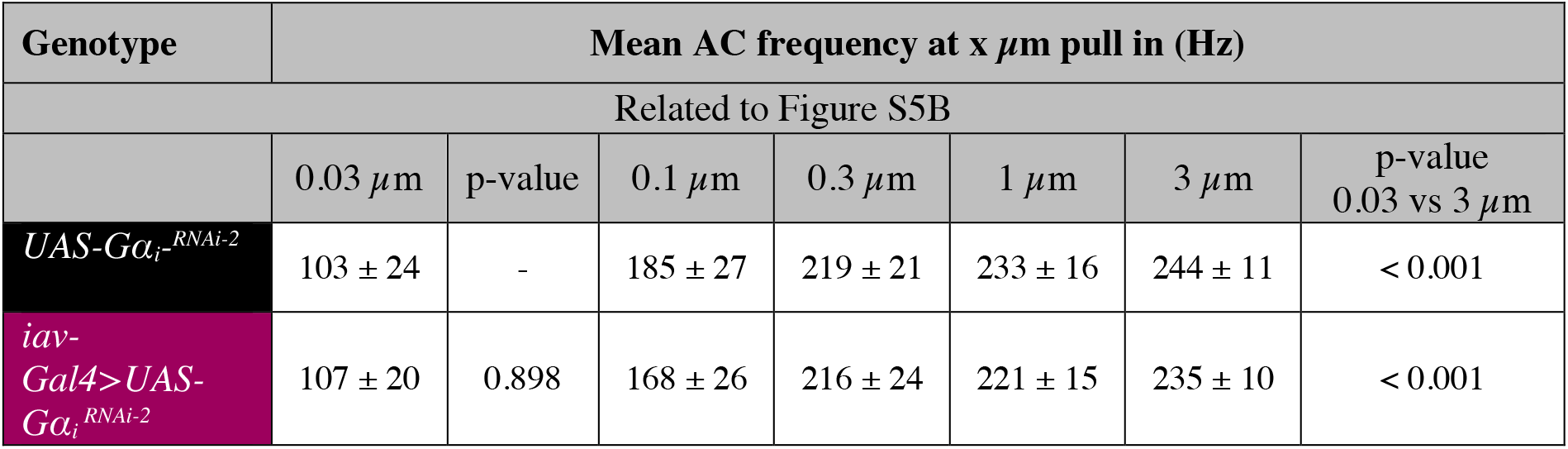
Mean AC frequency at different pull lengths in a second Gα_i_ protein knock-down. Quantification of AC frequencies of lch5 neurons at different pull lengths (0.03-3 μm). Values represent the mean ± SEM response of 10 lch5 organs from 10 individual animals. AC frequencies of both genotypes at 0.03 μm pulls were compared using an unpaired t-test (third column). AC frequencies at 0.03 and 3 μm in the same genotype were compared using paired t-tests (last column).

| Genotype | Mean AC frequency at x $\mu\text{m}$ pull in (Hz) | | | | | | |
| --- | --- | --- | --- | --- | --- | --- | --- |
| Related to Figure S5B |  |  |  |  |  |  |  |
| | 0.03 $\mu\text{m}$ | p-value | 0.1 $\mu\text{m}$ | 0.3 $\mu\text{m}$ | 1 $\mu\text{m}$ | 3 $\mu\text{m}$ | p-value<br>0.03 vs 3 $\mu\text{m}$ |
| *UAS-G$\alpha_i$<sup>RNAi-2</sup>* | 103 ± 24 | - | 185 ± 27 | 219 ± 21 | 233 ± 16 | 244 ± 11 | < 0.001 |
| *iav-Gal4>UAS-G$\alpha_i$<sup>RNAi-2</sup>* | 107 ± 20 | 0.898 | 168 ± 26 | 216 ± 24 | 221 ± 15 | 235 ± 10 | < 0.001 |

**Table S15.** Cell-specific Gα_o_ rescue. Quantification of AC frequencies of lch5 neurons at different pull lengths (0.03-3 μm). Values represent the mean ± SEM response of 10 lch5 organs from 10 individual animals. Unless stated otherwise (*), AC frequencies at 0.03 and 3 μm in the same genotype were compared using paired t-tests.

| Genotype | Mean AC frequency at x $\mu\text{m}$ pull in (Hz) | | | | | |
| --- | --- | --- | --- | --- | --- | --- |
| Related to Figures 6G + S5C |  |  |  |  |  |  |
| | 0.03 $\mu\text{m}$ | 0.1 $\mu\text{m}$ | 0.3 $\mu\text{m}$ | 1 $\mu\text{m}$ | 3 $\mu\text{m}$ | p-value<br>0.03 vs 3 $\mu\text{m}$ |
| *;; iav-Gal4* | 109 ± 14 | 198 ± 20 | 226 ± 16 | 224 ± 16 | 240 ± 15 | < 0.001 |
| *;Cirl<sup>KO</sup>; iav-Gal4* | 225 ± 16 | 227 ± 14 | 229 ± 13 | 226 ± 11 | 234 ± 11 | 0.609 |
| *;Cirl<sup>KO</sup>, UAS-G$\alpha_o$;* | 199 ± 21 | 220 ± 15 | 236 ± 18 | 238 ± 12 | 245 ± 8 | 0.074* |
| *Cirl<sup>KO</sup>, UAS-G$\alpha_o$; iav-Gal4* | 127 ± 18 | 190 ± 28 | 210 ± 18 | 237 ± 11 | 244 ± 9 | < 0.001 |
| *Wilcoxon Signed Rank test |  |  |  |  |  |  |

**Table S16.** Cell-specific Gα_o_ rescue. Comparison of AC frequencies generated at 0.03 μm pull. Values represent mean responses of 10 lch5 organs from 10 individual animals. All groups were compared using unpaired t-tests.

| Genotype | Comparison of mean AC frequencies at 0.03 $\mu\text{m}$ pull | | | | |
| --- | --- | --- | --- | --- | --- |
| Related to Figures 6G + S5C |  |  |  |  |  |
|  |  | *;; iav-Gal4* | *Cirl<sup>KO</sup>; iav-Gal4* | *;Cirl<sup>KO</sup>, UAS-G$\alpha_o$;* | *;Cirl<sup>KO</sup>, UAS-G$\alpha_o$; iav-Gal4* |
|  |  | 109 Hz | 225 Hz | 199 Hz | 127 Hz |
| *;; iav-Gal4* | 109 Hz | - | < 0.001 | 0.003 | 0.454 |
| *;Cirl<sup>KO</sup>; iav-Gal4* | 225 Hz | - | - | 0.339 | < 0.001 |
| *;Cirl<sup>KO</sup>, UAS-G$\alpha_o$;* | 199 Hz | - | - | - | 0.018 |
| *;Cirl<sup>KO</sup>, UAS-G$\alpha_o$; iav-Gal4* | 127 Hz | - | - | - | - |

